# Sequence heterochrony led to a gain of functionality in an immature stage of the central complex: a fly-beetle insight

**DOI:** 10.1101/2019.12.20.883900

**Authors:** Max S. Farnworth, Kolja N. Eckermann, Gregor Bucher

## Abstract

Animal behavior is guided by the brain. Therefore, adaptations of brain structure and function are essential for animal survival, and each species differs in such adaptations. The brain of one individual may even differ between life stages, for instance as adaptation to the divergent needs of larval and adult life of holometabolous insects. All such differences emerge during development but the cellular mechanisms behind the diversification of brains between taxa and life stages remain enigmatic. In this study, we investigated holometabolous insects, where larvae differ dramatically from the adult in both behavior and morphology. As consequence, the central complex, mainly responsible for spatial orientation, is conserved between species at the adult stage, but differs between larvae and adults as well as between larvae of different taxa. We used genome editing and established transgenic lines to visualize cells expressing the conserved transcription factor *retinal homeobox,* thereby marking homologous *genetic neural lineages* in both the fly *Drosophila melanogaster* and the beetle *Tribolium castaneum*. This approach allowed us for the first time to compare the development of homologous neural cells between taxa from embryo to the adult. We found complex heterochronic changes including shifts of developmental events between embryonic and pupal stages. Further, we provide, to our knowledge, the first example of *sequence heterochrony* in brain development, where certain developmental steps changed their position within the ontogenetic progression. We show that through this *sequence heterochrony*, an immature developmental stage of the central complex gains functionality in *Tribolium* larvae. We discuss the bearing of our results on the evolution of holometabolous larval central complexes by regression to a form present in an ancestor.

## Introduction

The brain is among the most complex organs of an animal, where sensory inputs and internal states are processed to guide its behavior. Hence, modifications of brain structure and function in response to specific requirements imposed by different life strategies and environmental conditions is paramount for each species’ adaptation. Insects represent one of the most diverse animal clades and they have conquered almost every habitat on earth (1–3). Indeed, based on the highly conserved basic bauplan within the insect clade, brains have diversified significantly in size, shape and position of their functional brain units, the neuropils (4–9). For instance, the mushroom bodies required for olfactory learning and memory are enlarged in bees, antennal lobes are reduced in aquatic beetles and the size of the optic lobes is increased in species that navigate in complex environments (9,13–17). In holometabolous insects, where larval stages often differ from the adult in life strategy and habitat, evolutionary adaptation imposes different brain morphologies even on successive life stages of one individual (18–20). Divergent brain morphologies emerge during embryonic and postembryonic ontogeny and, hence, any evolutionary modification depends on a modification of developmental mechanisms. Basic developmental processes appear to be conserved, reflecting the conserved basic architecture of the brain. Homology of neuroblasts and the resulting neurons is assumed (21–23) such that neuroblasts form conserved lineages. Based on this conserved process, evolution of developmental mechanisms is expected to act rather on details like the number of daugther cells formed, truncation of development and modification of lineage parts.

The low number of neural cells in insect brains compared to e.g. vertebrates, its basis of conserved lineages building up the brain, together with their experimental accessibility, makes insects an excellent choice to study the mechanisms of brain diversification during development. Despite the brain’s central role in insect evolution and the clade’s suitability to uncover underlying patterns, developmental mechanisms of brain diversification remain poorly studied.

Recent technical advances open the possibility to study such modifications of developmental mechanisms, both, in the classic model organism *Drosophila melanogaster* in order to pioneer the conceptual framework of neural development, and in other insects in order to reveal conserved and divergent aspects. The red flour beetle *Tribolium castaneum* is spearheading comparative functional work in neurogenesis due to its well-developed genetic toolkit and recent advances in neurobiological methods (24–38). Hence, establishing similar tools in *Drosophila* and *Tribolium* helps unravelling the developmental mechanisms of insect brain evolution through comparative developmental studies.

An intriguing evolutionary divergence in morphology of the developing brain – and therefore a suitable target for a *Drosophila – Tribolium* comparison – was found with respect to the central complex. The central complex is a neuropil that integrates multisensory information and acts predominantly as spatial orientation and locomotor control center (39–41). Related neuropils have been found in Crustaceans and Myriapods and it has even been further homologized to the vertebrate basal ganglia, while the homology of the central complex to the arcuate body of spiders is still discussed (42–46). In adult insects, the central complex is highly conserved consisting of a set of midline-spanning neuropils, the protocerebral bridge (PB), the central body (CB) consisting of an upper (CBU, or fan-shaped body, FB) and lower division (CBL, or ellipsoid body, EB) and the noduli (NO) with stereotypical patterns of innervation (Fig. 1A) (40,47–49).

**Fig. 1:**
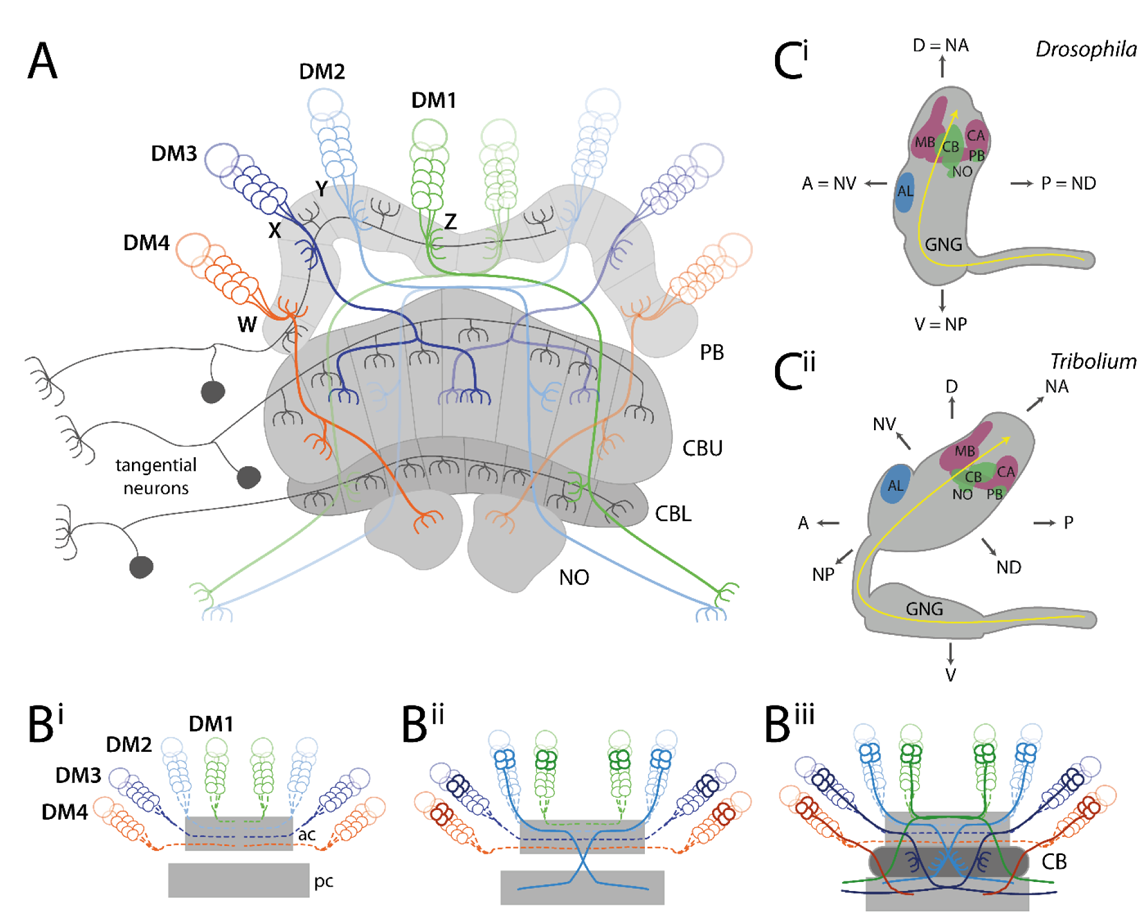
Structure and development of the central complex, and relationship of neuraxis to body axes. (A) Tangential neurons (dark grey) connect neuropils of the central complex with other areas. Columnar neurons (coloured) connect the different neuropils of the central complex with each other. Nearly all columnar neurons derive from four type II neuroblasts, DM1-4 (green, light-blue, dark-blue, orange) that project through WXYZ tracts. (**B)** Central complex development starts with the neurons of the DM1-4 lineage projecting into an anterior commissure (ac) (hatched lines in B^i^) where they cross the midline and build up a stack of parallel fibers. Later-born neurons (solid lines in B^ii^) undergo fascicle switching, i.e. they leave the fascicle at stereotypical locations and re-enter a fascicle of a posterior commissure (pc) forming X-shaped crossings with neurons from the contralateral side (called decussations) (B^ii^). Decussations occur at different points subdividing the future central body into columns (B^iii^). PB omitted for simplicity; based on (75,80,81). (**C)** The *Drosophila* (C^i^) and *Tribolium* (C^ii^) brains differ in their orientation within the head (lateral views). While the *Drosophila* brain is oriented perpendicular to the ventral nerve cord, the *Tribolium* brain is tilted backwards. This leads to discrepancies when using the body axis as reference. For instance, the AL is anterior in *Drosophila*, while it is more dorsal in *Tribolium*. Similarly, the PB is posterior in *Drosophila* but rather ventral in *Tribolium*. To facilitate cross-species comparisons, we use the neuraxis nomenclature as suggested by (82). In this system, the AL are n-ventral (NV) and the PB n-dorsal in both species. Shapes of brains are based on v2.virtualflybrain.org/ and data from this study, while the shape of the *Tribolium* GNG is from (83). Information about cell innervation in A was taken from (72,76,84). Abbreviations: AL antennal lobes, PB protocerebral bridge, CB central body, CBU upper division of the CB, CBL lower division of the CB, NO noduli, MB mushroom body (excluding CA), CA calyx, n neuraxis-referring, D dorsal, A anterior, V ventral, P posterior, GNG gnathal ganglia, DM dorso-medial, ac anterior commissure, pc posterior commissure.

In hemimetabolous insects, all neuropils develop during embryogenesis and already the hatchling has an adult-like central complex (50–52). By contrast, in holometabolous insects, the central complex forms partly during embryogenesis and is completed only during metamorphosis. In the tenebrionid beetles *Tenebrio molitor* and *Tribolium castaneum*, for instance, the larval central body consists of only one division, which was suggested to represent the upper division. The lower division was proposed to develop later during pupal stages (20,33,53). In *Drosophila*, no functional central complex neuropils are detectable in the first instar larvae. At that stage, the central complex anlagen consist of commissural tracts lacking neuropil morphology and characteristics of functionality, i.e. synapses and neuromodulator presence (54, 55). Only during late larval stages and metamorphosis, the central complex matures into the adult form (55–57). This divergent emergence of the CBU in different species is thought to correlate with the development of walking legs while the presence of the CBL may be linked to the formation of complex eyes (7,51,52,58). Intriguingly, the development at least of the upper division appears to be quite similar between the hemimetabolan desert locust *Schistocerca gregaria* and the fly *Drosophila melanogaster* albeit similar developmental steps occur at different stages (59).

This phenomenon represents a case of heterochrony (60, 61). Different definitions of this term have been proposed (62, 63). We use the term heterochrony to describe a change in developmental timing of a process in one taxon compared to other taxa. Such differences can be found with respect to development of shape, size and the time of maturation (60, 64) but also changes in the order of events within a developmental sequence can be interpreted in the framework of heterochrony (sequence heterochrony) (65–67). Heterochrony has strong influence on evolution: For instance, the accelerated frequency of somite formation in snakes contributes to their increased number of segments (68). Another example is the heterochronic extension of the growth phase of the postnatal infant human brain compared to other primates, leading to a relative increase of final brain size (60,69–71). The influence of heterochrony on insect brain evolution has not been thoroughly studied. Specifically, the observation of heterochrony in the central complex lacks detail because it is based on overall neuropil shape at two stages rather than a thorough comparison throughout development.

The central complex of the adult insect consists mainly of columnar and tangential neurons (49, 52). Tangential neurons connect other brain areas with one central complex neuropil (Fig. 1A) (52,72–74). In contrast, columnar neurons connect the different neuropils of the central complex with each other by projecting as four prominent tracts (the WXYZ tracts) from the protocerebral bridge into CBU, CBL, noduli and other brain structures (Fig. 1A) (47,49,75–78). These neurons are required for the formation of the typical columnar architecture of the central body and protocerebral bridge (47,52,75,79).

Important work in *Schistocerca gregaria* and *Drosophila melanogaster* revealed how such a complex innervation architecture is achieved during development (21,50,54,55,59,85,86). Specifically, the development of columnar neurons has been studied in detail: They stem from four neural lineages per hemisphere (Fig. 1A-B), called DM1-4 (alternative names in *Drosophila*: DPMm1, DPMpm1, DPMpm2, CM4 or in *Schistocerca*: ZYXW) (79,87,88). The respective neural stem cells (neuroblasts) are situated in the anterior-median brain close to the protocerebral bridge between brain hemispheres, i.e. in the pars intercerebralis. These lineages are built by type II neuroblasts, which generate approximately four times more cells than type I lineages (55,80,87,89,90). The neurites of these lineages first project ipsilaterally through the WXYZ tracts from the protocerebral bridge to the central body, where they turn and cross the midline forming a stack of parallel fibers (Fig. 1B^i^). Subsequent neurites leave the fascicle (de-fasciculation) and enter another fascicle of the brain commissure (re-fasciculation) to continue their growth to the other side of the brain, a process referred to as fascicle switching (81, 91). This happens at several stereotypical points along the commissure and symmetrically on both sides, such that neurites cross each other forming X-shaped crossings, which are called decussations (see (7) for distinction between ‘decussation’ and ‘chiasma’) (Fig. 1B^ii^-B^iii^). These decussations are the developmental basis for the typical columnar architecture particularly of the CBU (Fig 1B^iii^).

Studying such developmental processes of the central complex comparatively has been hampered by the lack of tools to mark homologous cells in two species. The elaborate toolkit of *Drosophila* for individual neural cell marking is not within reach in other organisms (92, 93) and even in *Drosophila* it has been challenging to mark neural lineages from embryonic neuroblast to the neurons of the adult brain. Recently, we suggested to compare homologous cells in different taxa by marking what we called *genetic neural lineages,* i.e. cells that express the same conserved transcription factor (33). Essentially, this approach assumes that transcription factors with conserved expression in the neuroectoderm and the brains of most bilateria are likely to mark homologous cells in closely related taxa throughout development. It should be noted, however, that the actual identity of a given neuroblast lineage is not determined by a single transcription factor but by a cocktail of several factors (94). Hence, genetic neural lineages may contain cells of several bona fide neural lineages. *Genetic neural lineages* can be labelled either by classic enhancer trapping, or a targeted genome editing approach, both available in *Tribolium* (28, 95).

In this study, we mark the *retinal homeobox (rx) genetic neural lineage* in both the red flour beetle *Tribolium castaneum* and the vinegar fly *Drosophila melanogaster* by antibodies and transgenic lines. We confirm the marking of homologous cells and subsequently scrutinize their embryonic and postembryonic development. We found a complex pattern of heterochrony underlying differentiation between larval and adult brains including the shift of certain developmental events between life stages. Intriguingly, we found that the order of developmental steps was changed, representing a case of *sequence heterochrony*, which to our knowledge had not been observed in the evolution of brain development before. As consequence, the larval central body of *Tribolium* represents an immature developmental stage, which gained functionality precociously. Apparently, central complex functionality does not require the full connectivity as observed in adult brains.

## Results

### Marking the *rx genetic neural lineage* in two species

To compare central complex development between two species, we wanted to mark a subset of homologous neurons that contribute to the central complex. For this purpose, we decided to use the *retinal homeobox (rx) genetic neural lineage* for three reasons: First, *rx* is one of the genes that is expressed almost exclusively in the anterior brain in bilaterians indicating a highly conserved function in many animals (96–104). Second, we had found projections into the central complex in a *Tribolium rx* enhancer trap line and a small subset of central complex projections in *Drosophila rx* VT-GAL4 lines (VDRC, # 220018, # 220016, discarded) (105, 106). Third, central complex phenotypes were observed in both *Drosophila* and *Tribolium* in a *Dm-rx* mutant and *Tc-rx* RNAi knock-down, respectively, indicating an essential role in central complex development (98, 107).

To mark *rx genetic neural lineages*, we first generated and validated an antibody binding the *Tribolium* Rx protein (Tc-Rx, TC009911) (Fig. S1) and used an available *Drosophila* Rx (Dm-Rx, CG10052) antibody (98). Next, we tested an enhancer trap in the *Tc-rx* locus (E01101; *Tc-rx-*EGFP line) (95) and confirmed co-expression of EGFP with Tc-Rx (Fig. S2). The enhancer trap marked a subset of 5-10 % of all Tc-Rx-positive cells in the adult and all EGFP-positive cells were Tc-Rx-positive as well (Fig. S2). For *Drosophila*, we generated an imaging line using CRISPR/Cas9 mediated homology-directed repair (Fig. S3). We replaced the stop codon of the endogenous *rx* locus with a P2A peptide sequence followed by an EGFP coding sequence (28,108,109). The resulting bicistronic mRNA led to translation of non-fused Dm-Rx and EGFP proteins (*Dm-rx-*EGFP; Fig. S3), and so, our analysis revealed complete co-expression of Dm-Rx and EGFP. Based on both antibodies and transgenic lines we tested the labelled cells for homology.

### Similar location of *rx*-positive neural cell groups in both species

To get an overview on the conservation of Rx expression between *Drosophila* and *Tribolium*, we first compared the location of Rx-positive cells in adult brains and embryos. Note that the axes of the brain relative to the body axes are not conserved in insects. Therefore, we describe the location according to the ‘neuraxis’ for both species, where ‘*Drosophila* posterior’ becomes neuraxis-dorsal (n-dorsal) while ‘*Drosophila* dorsal’ equals neuraxis-anterior (n-anterior) (see explanation in Fig. 1C). We found four major domains of Rx-positive cells (I-IV) located in similar regions in both species (Fig. 2A-B; see stacks and videos of all projections shown in this paper on figshare: figshare.com/account/home#/projects/64799). Specifically, cells of cluster IV surrounded the protocerebral bridge in both species in a pattern similar to DM1-4 lineages. In embryos of both species, Rx was expressed in the labrum (arrowheads in Fig. 2C-D) as well as in corresponding regions of the anterior-lateral part of the neuroectoderm (arrows in Fig. 2C-D).

**Fig. 2:**
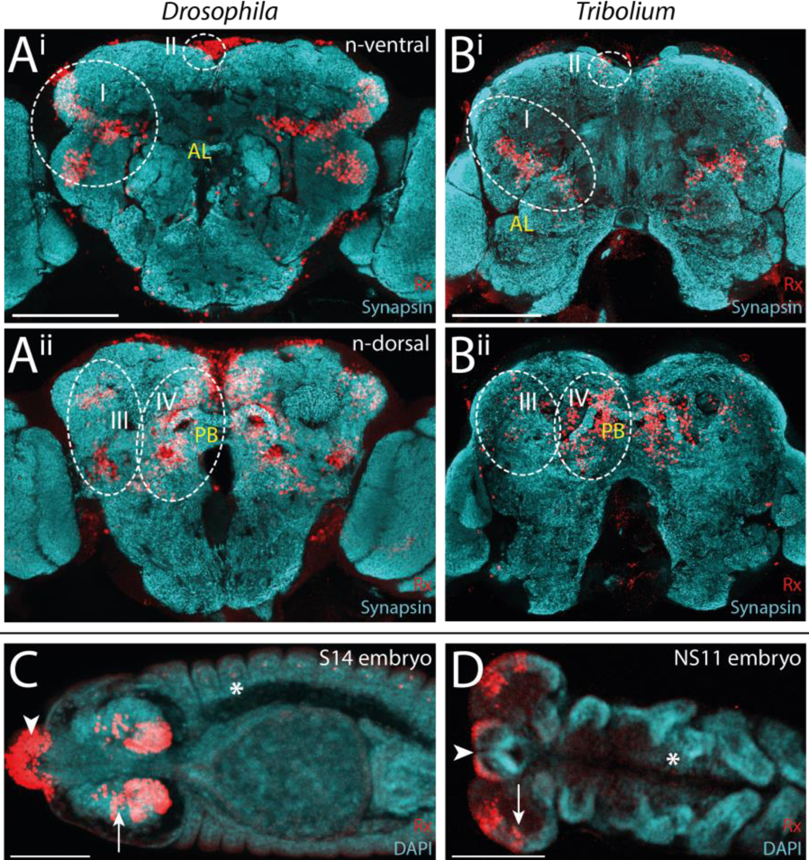
Rx expression is conserved in *Drosophila* and *Tribolium* adult brains and embryos. (A-B) Immunostainings against Rx and synapsin in both species revealed four domains of Rx-positive cells (I-IV, dotted white lines) with similar shape and position within the brain. Shown are n-ventral (i) and n-dorsal views (ii) (Fig. S4). **(C-D)** In *Drosophila* (S14) and *Tribolium* (NS11) embryos Rx was expressed in the labrum (arrowhead) and in similar regions of the lateral head neuroectoderm (arrows). In addition, single cells of the peripheral nervous system and ventral nerve cord were labelled in each segment (asterisk; Fig. S1). Note that the head lobes of *Tribolium* embryos are shown as flat preparations while the *Drosophila* head was imaged within the egg. According to the *bend and zipper model* of head morphogenesis, the expression patterns are very similar (110). Abbreviations like in Fig. 1. Scale bars represent 100 µm.

Next, we asked to which lineages these domains of the adult brain belong. To this end, we related Rx-positive cell groups to maps of neural lineages of the *Drosophila* brain (111, 112) and tentatively transferred the nomenclature to *Tribolium* (Fig. S4). For this, we also included prominent projections of Rx-positive cells marked by the transgenic lines, to substantiate our assignments (Figs. S2-4). Rx-positive cell groups likely belonged to eleven neural lineages (Fig. S4, Table S1 and Supporting Results). Four of these (DM1-4) were prominently marked in the imaging lines of both species. Because DM1-4 are known to contribute to the central complex we focused on the comparison of Rx-positive cell clusters of these lineages.

### Central complex Rx-positive cell clusters are homologous between *Drosophila* and *Tribolium*

To corroborate the homology of Rx-positive DM1-4 neurons, we examined the location and projection pattern of these cell clusters in detail. We indeed found similar cell body locations around the protocerebral bridge (Fig. 3A-B) and similar projection patterns into the CBU (Fig. 3C-D), CBL and noduli (Fig. 3E-F) in both species. The similarity relative to central complex neuropils was visualized in 3D reconstructions (Fig. 3G-H, see videos on Figshare) and allowed us to define homologous cell clusters. Given the lack of a detailed map and homology assessments for the *Tribolium* brain, we assigned the fiber bundles MEF, dlrCBU and mrCBU (dlrFB, mrFB, see e.g. (54)) based on their similarity to the *Drosophila* brain and the novel lineage information gained in this study (Fig. S4, Supporting Results). Specifically, in both species, DM4 cell bodies lay around the lateral tip of the protocerebral bridge and their axons projected through the medial equatorial fascicle (MEF) (orange in Fig. 3). Cell bodies of DM2/3 (light and dark blue, respectively, in Fig. 3) were close to each other at the n-anterior bend of the protocerebral bridge. Their axons projected into distinct tracts through the dorsal root of the CBU (dlrCBU). DM1 cell bodies (green in Fig. 3) lay near the midline and their axons projected through the medial root of the CBU (mrCBU).

**Fig. 3:**
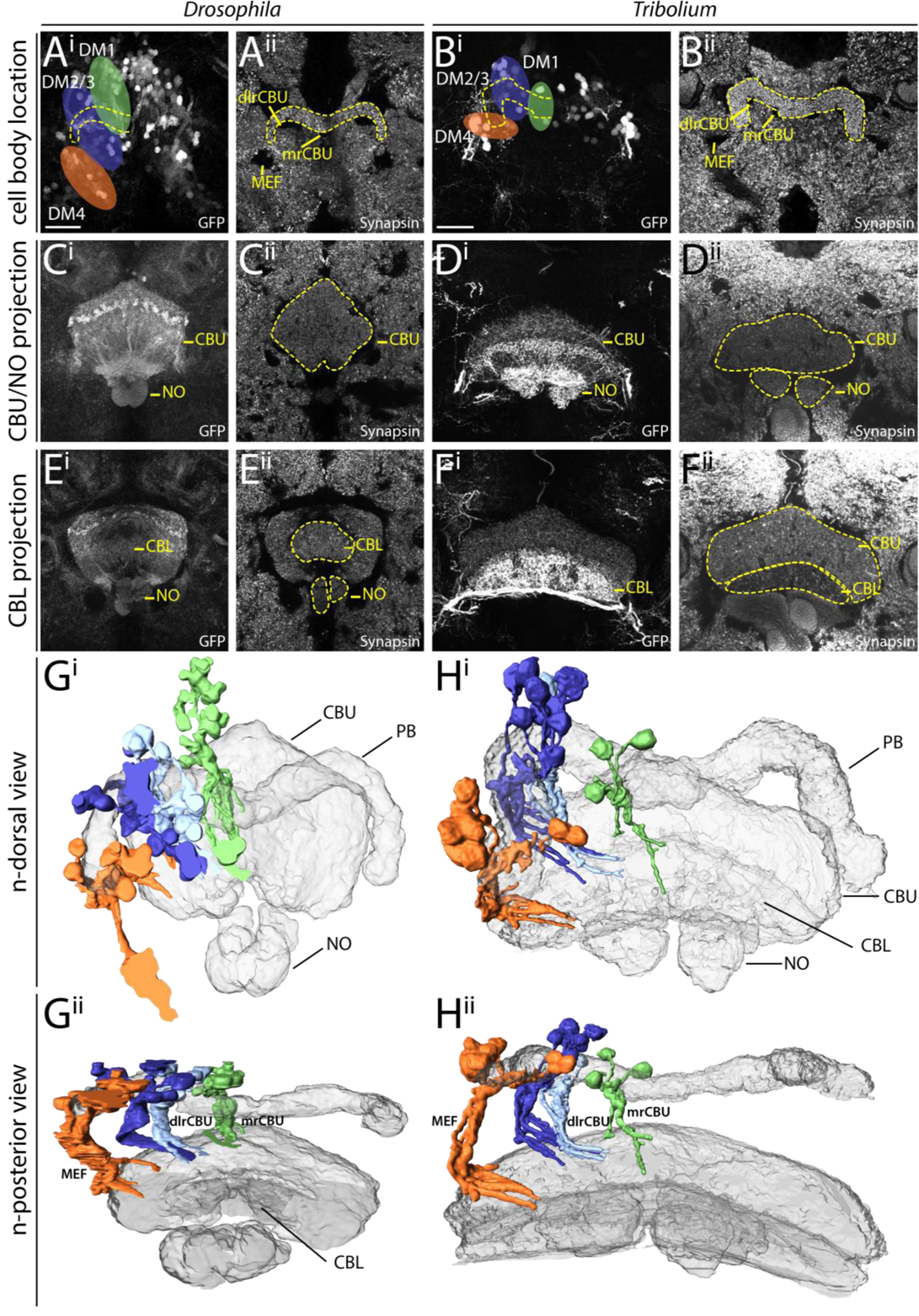
Homologous Rx cell clusters contribute to the adult central complex columnar neurons of lineages DM1-4. A to F depict substacks of *Drosophila* (left columns) and *Tribolium* (right columns) adult brains on which the 3D reconstructions in G and H are based. Homology of cell clusters and DM1-4 lineages was based on three criteria: Rx expression (i.e. being part of the *rx genetic neural lineage*), similar cell body location and similar projection patterns. **(A-B)** Cell groups of lineages DM1-4 (coloured areas) around the protocerebral bridge (yellow dotted lines) are shown for *Drosophila* (A) and *Tribolium* (B). **(C-D)** Projection pattern of GFP expressing neurites of these cell groups in the CBU and NO fraction. **(E-F)** Much less signal was found in the CBL fraction. Note that the *Tribolium* DM4 group had a very high GFP expression level, such that those projections were particularly visible in the CBL (see Figs. S2-3 for transgenic line information). **(G-H)** 3D reconstructions of synapsin staining (grey-transparent) and the EGFP marked cells of DM1-4 lineages. G^i^/H^i^ depicts the n-dorsal view shown in A-F. G^ii^/H^ii^ is rotated to an n-posterior view with the central complex coming to the front to judge similarity of the tract architecture. Similar stereotypical positions were found in both species for DM1 (green), DM2/3 (blue shades – sharing a fiber bundle) and DM4 (orange). Due to the large number of labelled cells within the CB, the projections could not be followed further. GFP channels (i) are maximum intensity projections, while synapsin channels (ii) are SMEs (113). Abbreviations as in Fig. 1. dlrCBU dorsal root of the CBU (synonym: dlrFB), mrCBU medial root of the CBU (mrFB), MEF medial equatorial fascicle. Scale bars represent 25 µm and apply to all panels of each species.

Our classification of these Rx expressing cell clusters to lineages DM1-4 was corroborated in *Drosophila* by Rx immunostainings in the R45F08-GAL4 line, a *pointed* GAL4 enhancer construct that was suggested to label a large subset of neurons of the DM1-3 and 6 lineages (55). Moreover, we crossed the *Dm-rx-*EGFP line to the R45F08-GAL4 line. We found that approximately 90 % of R45F08-GAL4 marked cells also expressed Rx (Fig. S5A-B). In addition, a substantial part of the midline projections overlapped between both transgenic lines (Fig. S5C).

Note that the *Dm-rx*-EGFP line marked all Dm-Rx-positive cells while the *Tc-rx*-EGFP line marked only a subset of Rx-positive cell bodies (Figs. S2 vs. S3). This resulted in more prominently marked tracts in *Drosophila* compared to *Tribolium*. However, during development, the number of EGFP-positive DM1-4 cells increased in *Tribolium* resulting in thicker projections (Fig. S2). Especially, the *Tribolium* DM4 Rx expressing group showed a very high EGFP expression, such that the respective projections into the CBU, noduli and CBL as well as the connections to the lateral accessory lobes appeared much stronger than in *Drosophila* (Fig. 3 B/D/F^i^). This divergence of intensity was likely a particularity of the *Tribolium* enhancer trap.

In summary, we assume homology of the Rx-positive cells of the DM1-4 lineages of *Drosophila* and *Tribolium* based on the shared expression of a highly conserved brain regulator and the specific similarity of cell body location of the DM1-4 lineages relative to the protocerebral bridge and their similar projection patterns in adult brains. Note that *rx* is expressed in most but probably not all cells of the DM1-4 lineages and in addition is expressed in cells contributing to other brain regions like the mushroom bodies, which were not examined here. The DM1-4 lineages are key components of the central complex, providing nearly all columnar neurons (54,59,114). Therefore, the *rx genetic neural lineage* is an excellent marker to compare central complex development between fly and beetle.

### Divergent central complex structures in the L1 larva of *Drosophila* and *Tribolium*

Next, we examined central complex structures in the first instar larval (L1) brain of both species, since the strongest divergence between *Drosophila* and *Tribolium* seemed to occur at the larval stage. Here, tenebrionid beetle larvae have a partial central complex neuropil already at the larval stage (20, 33) while in *Drosophila* L1 larvae any central complex neuropil is missing (54). Our imaging lines allowed us to compare DM1-4 innervation and resulting central complex structures at the L1 stage of both species complemented by synapsin and acetylated α-tubulin staining (115) to reveal functionality and underlying tract architecture, respectively (Fig. 4).

**Fig. 4:**
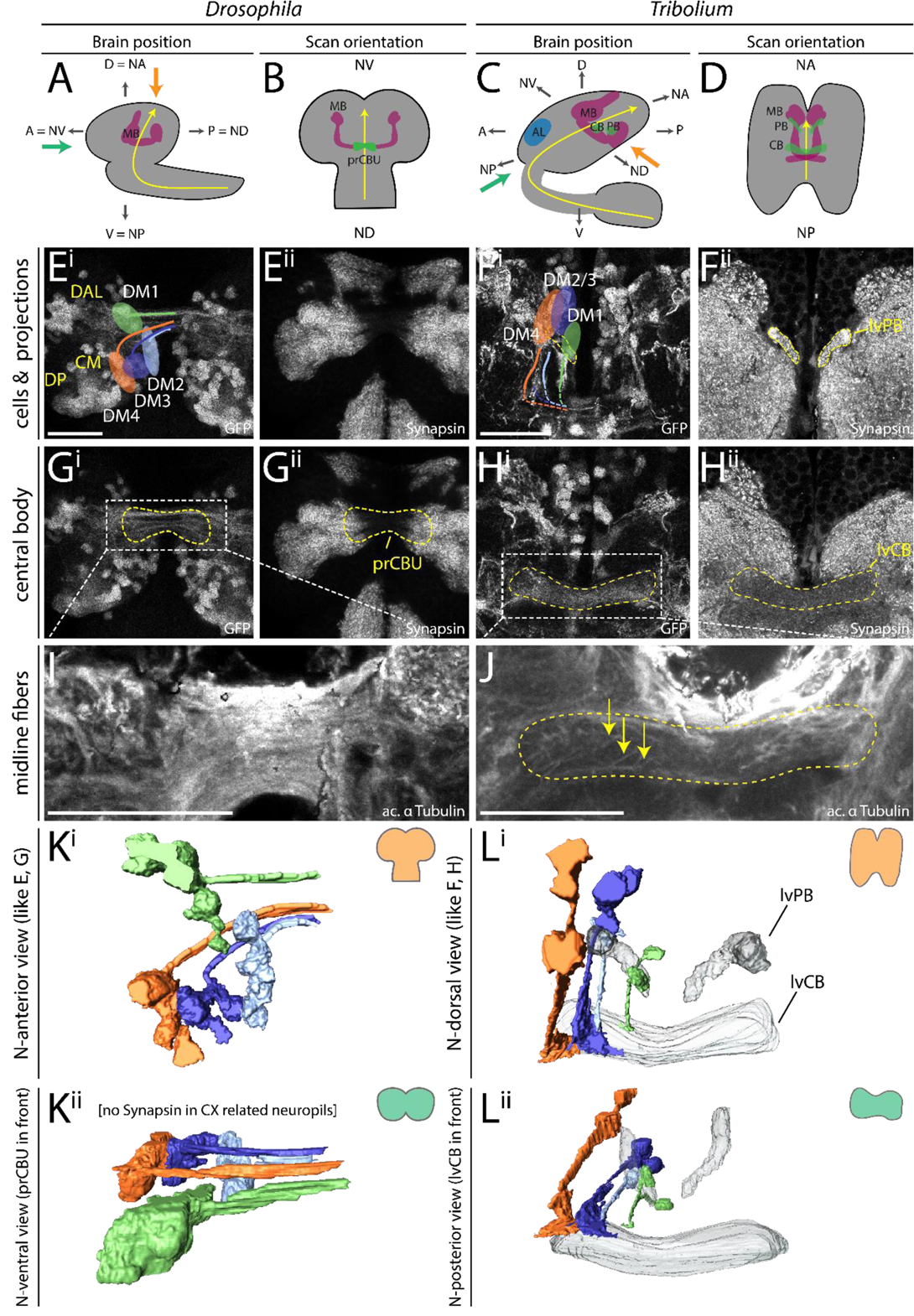
Different patterns of DM1-4 projection and central complex morphology at the first larval stage. (A+C) The *Drosophila* (left columns) and *Tribolium* (right columns) L1 brains are positioned differently within the head, visualized by lateral views in A and C. Indicated are the denominators for anterior, posterior, dorsal and ventral (A,P,D,V) for both body axes and neuraxes (with prefix N). Further shown are the curved neuraxis (yellow) and the larval neuropils MB (magenta), AL (blue), CB and PB (green). The orange arrows indicate the different directions of the performed scans. The green arrows indicate the orientation displayed in K/L^ii^ where central complex structures are best visible for both species. **(B+D)** The brains are depicted as they were scanned in E-J (i.e. from the angle of the orange arrow). **(E-H)** Differences between species were observed in cell cluster position and projection patterns as well as neuropil architecture. In *Tribolium*, arrangement and projection were already similar to the adult (compare L with Fig. 3) although the PB was still split. In *Drosophila* it differed dramatically: No central complex neuropils were detected and the DM1-4 lineages projected straight across the midline. In E^i^ the approximate position of other lineages of the *Drosophila* brain are shown, i.e. dorso-anterio-lateral (DAL), dorso-posterior (DP) and centro-medial (CM) lineages (yellow). **(I-J)** Anti-acetylated-α-Tubulin immunostaining revealed that in *Drosophila* midline-spanning fibers build up a simple stack of fascicles, containing the primordial central body. In *Tribolium,* in contrast, the functional central body contains already some decussated fibers. **(K-L)** 3D reconstructions visualize the spatial relationship between the lineages and highlight the differences between the species. Upper panels (i) reflect the orientation shown in E-H, while in the lower panels (ii) are oriented such that the prCBU and lvCBU are in front, i.e. the central complex is shown in a comparable perspective (see green arrow in A and C). Abbreviations like in previous figures. lv larval. Scale bars represent 25 µm.

The position of the brains within the L1 larva differs between the species, which has to be considered when comparing them (see scheme in Fig. 4A-D). As previously described (54), we found no functional (i.e. synapsin-positive) central complex neuropil in *Drosophila* L1 (neither protocerebral bridge, central body nor noduli; Fig. 4E^ii^,G^ii^). In *Tribolium*, in contrast, we observed a protocerebral bridge, which in synapsin stainings was non-fused (Fig. 4F^ii^). Further, we found a larval central body (lvCB), which showed no morphological sign of subdivision into upper or lower division (Fig. 4H^ii^). Moreover, neither neuropil displayed a columnar structure in anti-synapsin or anti-GFP stainings (Fig. 4F^ii^/H). Hence, the lvCB appeared as a simple bar-shaped neuropil.

The analysis of Rx expressing DM1-4 cells in *Drosophila* revealed that the spatial arrangement of marked cell bodies and their projections in the L1 differed from the adult (Fig. 4E^i^ – compare to Fig. 3A). The cell clusters differed both, in their position within the brain and with respect to each other. To correctly assign their identities to DM1-4 despite such divergence, we used their projections across the midline as hint and compared the location of the marked cell bodies with recent lineage classifications based on EM data (54). Most strikingly, the cell bodies of the DM2/3 lineages were not yet located between DM1 and DM4 (Fig. 4E^i^/K^i^). In *Tribolium*, in contrast, the DM1-4 cell clusters had an arrangement along the larval protocerebral bridge like the adult situation (Fig. 4F^i^/L^i^).

The projection patterns of the *rx*-positive DM1-4 lineages differed between the two species, as well. In *Drosophila,* they formed a straight common projection across the midline in a bundle of parallel fascicles as described before (Fig. 4E^i^/G^i^/I) (54). Decussation was not found, neither on the level of EGFP signal nor based on acetylated α-tubulin staining (Fig. 4G^i^/I). In *Tribolium*, in contrast, the neurites projected first parallel to the midline towards n-posterior, projecting through (in the case of DM1-3) or passing by the protocerebral bridge (DM4). Then, they described a sharp turn towards the midline projecting medially into the lvCB neuropil towards the other side (Fig. 4F/H/L). Basically, this pattern resembled the adult one (compare Fig. 4L^i^ with 3H). In contrast to *Drosophila* L1, acetylated α-tubulin staining (but not EGFP signal) revealed a system of crossing, i.e. decussated fascicles in the region of the lvCB (Fig 4J). This pattern of decussation differs strongly from that found in the pupa in that it is built by less prominent fascicles and not visible with the *Tc-rx*-EGFP line (Fig. 9).

In summary, we confirm that *Tribolium* but not *Drosophila* has a functional (i.e. synapsin-positive) central body and protocerebral bridge at the L1 stage. We further show that the DM1-4 lineages of *Tribolium* larvae already ressemble the adult pattern including some decussations, while this is not the case in *Drosophila*. We note that, despite the presence of four adult-like WXYZ tracts and first decussations in the *Tribolium* L1, the lvCB is not visibly divided into columns. This contrasts with *Drosophila* where the presence of adult-like tracts and the start of decussation coincides with the division into columns in the early pupa (Fig. 8). Hence, heterochrony is found with respect to the gain of functionality at the L1 stage and with respect to the development of the underlying neural lineages.

Importantly, the functional *Tribolium* larval central body did not represent an adult-like upper division. Rather, it morphologically corresponded to a developmental step found in other species. Specifically, the initiating decussations within a tract of largely parallel fibers mirrors the situation seen in an embryonic stage of the grasshopper (59, 114). Hence, the *Tribolium* larval central body represents a case of heterochronic gain of functionality of an immature developmental stage rather than a heterochronic shift of the development of an adult-like structure.

### Embryonic central complex development proceeds faster in *Drosophila*

We next asked whether the observed differences were explained by simple temporal shifts within a conserved developmental series or whether certain steps changed their position in the series (i.e. sequence heterochrony). For this we compared discrete developmental events of the central complex in both *Tribolium* and *Drosophila*. We made use of our *rx* imaging lines to compare the development of homologous cells by defining three events identifiable in embryos of both species. These were the first axon projection emerging from marked cells, the first midline-crossing projection and the stage, when a larva-like projection pattern was reached. Further, the emergence of functional central body and protocerebral bridge as judged by synapsin staining was examined. Given the large differences in absolute developmental time between *Tribolium* and *Drosophila* we used relative developmental time.

The first axons of the *rx genetic neural lineages* formed at a similar relative timing in both species (*Drosophila* 37 % developmental time, *Tribolium* 39 %; Fig. 5A-B, see Material and Methods and Supporting Information for all staging details). The appearance of the first midline-crossing projection appeared earlier in *Drosophila* than in *Tribolium* (*Drosophila* 43 %, *Tribolium* 58 %; Fig. 5C-D). Likewise, the ‘final’ larval-like pattern was reached much earlier in *Drosophila* (51 %, *Tribolium* 81 %; Fig. 5E-F). However, at this stage the tracts of DM1-4 in *Tribolium* were already in similar spatial orientation and projection pattern as found in the adult. Moreover, despite this slower pace of development, *Tribolium* performed two more steps during embryogenesis, which in *Drosophila* were postembryonic: We found weak decussations and gain of functionality in the prospective central body region (i.e. synapsin staining), in late stage *Tribolium* embryos at approximately 81% (Fig. 5G-H). A distinct protocerebral bridge or central body that was clearly differentiated from other areas was not detectable in the embryo, however.

**Fig. 5:**
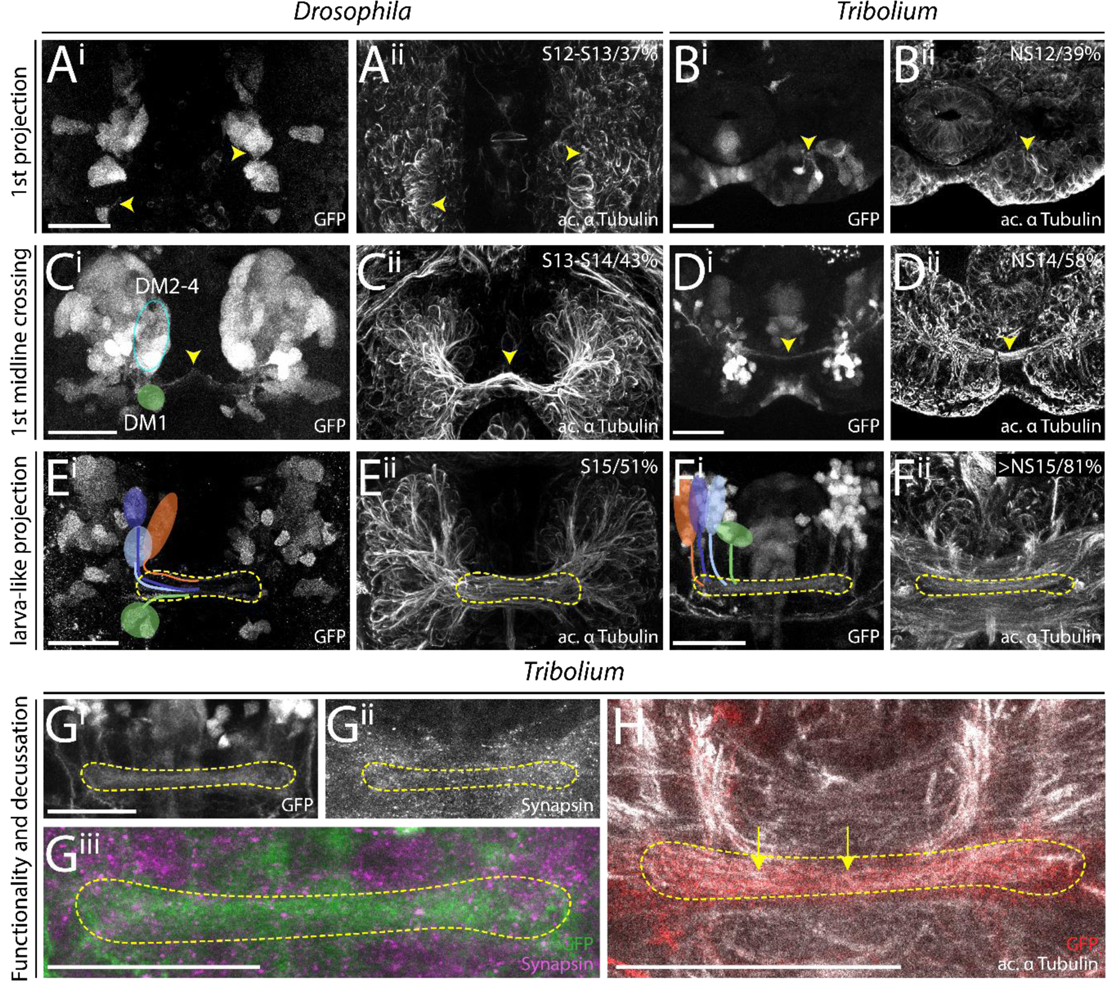
Key events of central complex development occur during late embryogenesis in *Tribolium* but not *Drosophila*. Comparable steps of central complex development are shown in one row for both species, the respective stage/relative time of development are shown in panels ii. The analysis was based on EGFP-labelled neurons (panels i) while acetylated α-tubulin staining is shown for reference (panels ii). **(A-B)** The development of the first axons happened at a similar time in *Drosophila* and *Tribolium*. **(C-D)** First midline-crossing fibers appeared earlier in *Drosophila*. **(E-F)** Likewise, the larva-like projection pattern was reached earlier in *Drosophila*. **(G-H)** The late-stage embryonic central complex of *Tribolium* is already faintly synapsin-positive (G^ii^, magenta in G^iii^) while the *Drosophila* lvCB remains synapsin-negative (not shown). In *Tribolium*, first decussations were visible as well (H, yellow arrows). Note that the assignment of Rx-positive cell clusters to the DM1-4 lineage groups was not unambiguous before mid-embryogenesis. Tentatively, we indicated the location of DM1 (green) and DM2-4 cells (blue oval form) in C^i^. Later, the groups could be assigned to DM1-4 lineages (E-F). Stages in *Drosophila* correspond to (116) and in *Tribolium* to (25). Posterior is up, except in panels F, G and H where dorsal is up. Scale bars represent 25 µm.

We conclude that both species initiated development of the *rx genetic neural lineage* at a comparable time of development, that *Tribolium* proceeds slower but eventually includes two more developmental steps in embryogenesis. This represented a pronounced heterochronic shift of conserved developmental steps between different life stages. More strikingly, certain steps of the developmental series switched their order representing a case of *sequence heterochrony* in brain diversification (Fig. 6). Specifically, the decussation and an adult-like tract organisation occurred before the larval growth phase of the lvCB in *Tribolium* but after that stage in *Drosophila*.

**Fig. 6:**
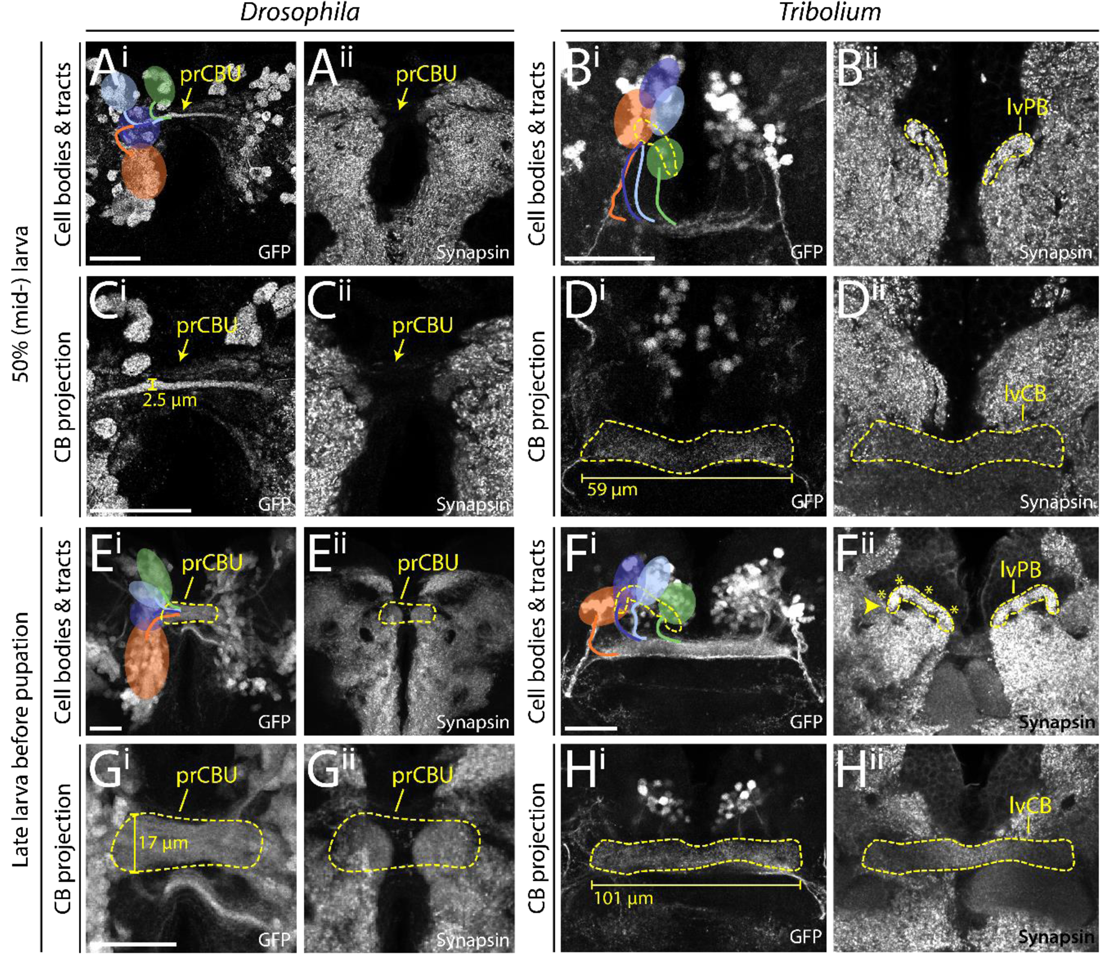
In both species, the *rx genetic neural lineage* shows substantial growth. During larval stages the identified cell clusters and their projections retained their position, but proliferated so that larger cell clusters and thicker and larger projections were built. **(A-D)** Depicted are projections at mid-larval stages (50 % of larval developmental time) where cell number and projections have qualitatively increased in number and size, respectively. **(E-H)** Shown are late larval stages before pupation, where cell numbers and projection sizes have increased greatly from 50 %. The late lvPB of *Tribolium* can be divided into discrete columns already, indicated by four asterisks on one hemisphere. Bars in C, D, G and H indicate the size increase of midline structures. In *Drosophila*, the prCBU increased in width from 2.5 to 17 µm from 50 to 95 % of larval development. In L1, the prCBU is non-distinguishable using the *rx*-GFP line. The central body of the *Tribolium* L1 brain displayed in Fig. 4 was 51.6 µm long, the mid-larval lvCB was 58.7 µm and the late larval lvCB was 100.9 µm long. For *Drosophila* n-ventral and for *Tribolium* n-anterior is up (see Fig. 4 for details). Abbreviations like in previous figures; pr primordium. Scale bars represent 25 µm and apply to panels i and ii and in case of *Tribolium* to D and H, respectively.

### In the larva, central complex structures grow but do not change basic morphology

Next, we asked how central complex structures changed during the larval period from the starting L1 architecture. We examined the position of cell clusters and their projections at 50 % (Fig. 6A-D) and at the end of the larval period (∼ 95 %) (Fig. 6E-H). In *Drosophila*, the primordium of the CBU increased in thickness, particularly after 50 % of larval development (compare Fig. 6C^i^ to G^i^) but it remained devoid of synapsin (Fig. 6C^ii^/G^ii^). In line with the literature we detected no decussations during larval stages. However, the position of DM1-4 cell clusters changed in *Drosophila*. Until 50 % of larval development, DM2 and DM3 cell bodies shifted n-ventrally, taking a position between DM1 and DM4 (compare Fig. 4E with Fig. 6A^i^). Towards the end of larval development, cell clusters became arranged in a straight line along the neuraxis, DM1 most n-ventral, DM4 most n-dorsal (Fig. 6E^i^).

In *Tribolium*, the central body grew in length and thickness as well (compare Fig. 6D^i^ with H^i^). In addition, the position and shape of the protocerebral bridge changed. In L1 and 50 % larval brains, the separate parts of the protocerebral bridge were still oriented along the n-anterior/posterior axis. In late larval brains, however, they shifted into a position more perpendicular to the neuraxis. Accordingly, the positions of the marked cell clusters remained constant in the first half of larval development (Fig. 6B^i^/H^i^). However, from 50 % to the end of the larval period they became arranged in one line along the protocerebral bridge. We also identified the presence of a columnar architecture in the protocerebral bridge in the late larva as judged by anti-synapsin staining, hence, there were four distinguishable sub-areas per hemisphere, likely reflecting a localized innervation near each of the four tracts. In both species’ larval brains, we qualitatively observed an increase in cell number of the DM1-4 *rx*-positive cell bodies.

We conclude that the larval period of central complex development is characterised mainly by growth of the central complex neuropils in both species. Apart from some shifts of cell body location, the structure established during embryogenesis was mostly maintained during the larval period. Importantly, the *Drosophila* central complex precursor remained synapsin-negative while in *Tribolium* both the lvCB and lvPB remained synapsin-positive, thus still resembling an immature structure throughout the larval period.

### The *Drosophila* central complex acquires functionality at later stages of pupal development

Last, we examined pupal stages to reveal when heterochronic divergence in early central complex development was eventually levelled out to reach the conserved adult structure. We stained brains of 0 (prepupal stage), 5, 15, 20, 30 and 50 % of pupal development (see Supporting Material and Methods for staging) for EGFP and synapsin. In *Drosophila,* the protocerebral bridge appeared at 5 % of pupal development (Fig. 7C^i^), grew subsequently and fused medially between 30 and 50 % of pupation (Fig. 7I/K^i^). Columns became visible at 15 % (Fig. 7E^i^). The upper division of the *Drosophila* central body appeared first at 5 % of pupal development (Fig. 7C^ii^). Strength of synapsin staining increased at 15 %, coinciding with the emergence of layers and columns structuring the CBU (arrows and bars, respectively, Fig. 7E^ii^). This coincided with *Dm-rx-*EGFP projections forming a columnar division (Fig. 8C^iii^). Thickness increased from 30 % onwards resulting in the fan-like structure typical for the *Drosophila* CBU (Fig. 7G/I/K^ii^). The *Drosophila* CBL emerged later at 15 % pupation, (Fig. 7E^iii^) and continued bending until it formed the typical toroid form that was nearly closed at 50 % pupation (Fig. 7K^iii^). Noduli appeared at the same time as the CBL as one paired subunit at 15 % of pupation (Fig. 7E^ii^), and only at 50 % an additional subunit was detected (Fig. 7K^ii^). Note that adult noduli are eventually comprised of three to six subunits, which apparently developed after 50 % development (78).

**Fig. 7:**
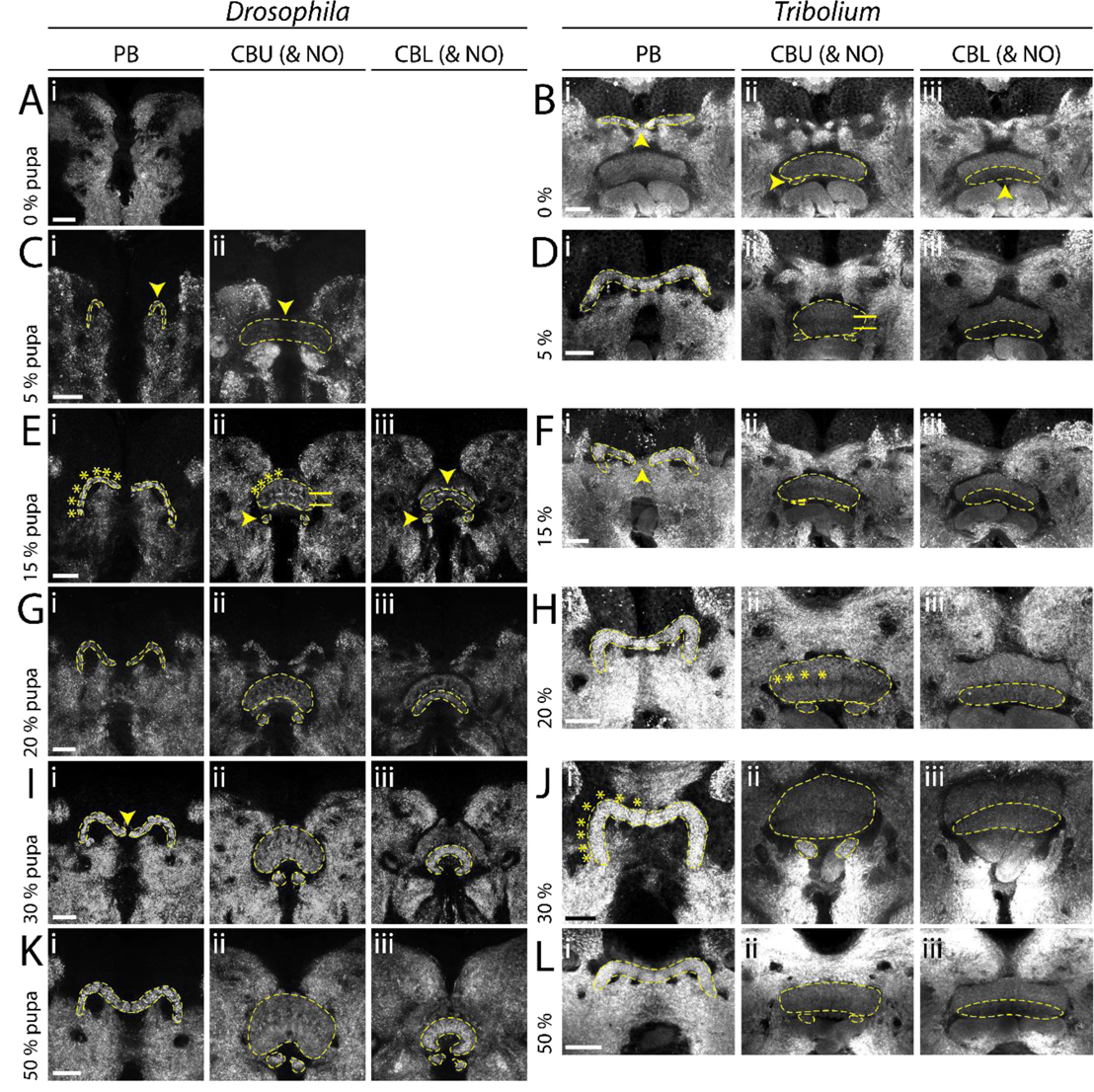
Pupal central complex development of *Drosophila* is delayed compared to *Tribolium*. Displayed are substack projections of an anti-synapsin staining of the same preparations used for tracing Rx-positive cell clusters in Figs. 8-9. **(A-D)** At 0-5 % pupation, in *Drosophila* the first functional neuropils have appeared, while in *Tribolium* NO and CBL have appeared and the CBU developed layers. **(E-H)** At 15-20 % pupation, the *Drosophila* central complex develops columns and layers, and NO and EB appear. In *Tribolium*, columns develop and the PB fuses. **(I-L)** At 30-50 % pupation, central complex structures resemble the adult, as the PB develops columns and fuses. Note that through slight deviations in positioning of the pupal brains, the CBU appears thicker in some stages than in others (e.g. H versus J). Following *Drosophila* events are highlighted by yellow arrowheads: Appearance of a functional PB (C^i^), CBU (C^ii^), NO (E^ii/iii^) and CBL (E^iii^), and last stage of an unfused PB (I^i^). Following *Tribolium* events are highlighted by yellow arrowheads: The last stage of an unfused PB (B^i^, F^i^, note the variability in the timing of fusion), appearance of NO (B^ii^) and CBL (B^iii^). A division into distinct layers in the CBU are marked by horizontal bars. A division into columns in the PB and CBU is marked by asterisks. Abbreviations like in previous figures. Scale bars represent 25 µm.

In *Tribolium*, the larval protocerebral bridge developed further and fused between 5 and 20 % (Fig. 7D/F/H^i^; note that we observed a higher heterogeneity in our *Tribolium* dataset with respect to protocerebral bridge fusion and other events). A division into eight columns typical for the adult neuropil became visible at 30 % on the level of synapses (Fig 7J^i^). However, based on the synapsin and EGFP signal of the *Tc-rx*-EGFP line, a division of the central body into columns was less visible at any developmental stage compared to *Drosophila*. This is in line with previous observations that central complex columnar architecture can be visible to quite different degrees in different taxa (12). Separate upper and lower divisions of the central body became visible already at the beginning of pupation (Fig. 7B^ii/iii^). The CBU increased in size, and at least two layers became visible at 5 % (Fig. 7D^ii^). The subdivision into columns was faintly visible from 20 % onwards (asterisks in Fig. 7H^ii^). The CBL appeared right at the beginning of pupation with weak synapsin signal intensity (Fig. 7B^iii^), which increased from 15 to 20 % of pupation (Fig. 7F/H^iii^). Noduli appeared at the prepupal stage (Fig. 7B^ii^). They thickened considerably at 20 % pupation (Fig. 7H^ii^) building two subunits between 30 and 50 % (Fig. 7J/L^ii^) eventually showing three subunits in the adult (not shown).

We concluded that protocerebral bridge, central body and noduli emerge later in the *Drosophila* pupal brain compared to *Tribolium*. Importantly, during pupation, the *Tribolium* larval central body matures significantly becoming quite different from its larval appearance corroborating that in larvae, the central complex is an immature but functional developmental stage.

### The *rx genetic neural lineages* contribute in a similar way to build the central complex during metamorphosis in both species

Given the overall heterochronic development of the central complex we asked in how far the development of the *rx genetic neural lineage* reflected these differences during metamorphosis.

In *Drosophila* pupal brains, the array of DM1-4 cell clusters turned from their straight orientation along the midline into a bent configuration following the protocerebral bridge (Fig. 8A-F^i^). The corresponding tracts underwent massive rearrangement, with typical bends similar to an adult configuration already visible at 5 % pupation. Most notably, decussations were created by fascicle switching of the DM1-3 tracts starting at 5 % of pupal development (Fig. 8B^ii^) and became prominent from 15 % onwards (Fig. 8C^ii^). This resulted in a parallel columnar organisation of the CBU at 15 % (Fig. 8C^iii^) and the marked tracts at 20 % (Fig. 8D^ii^).

**Fig. 8:**
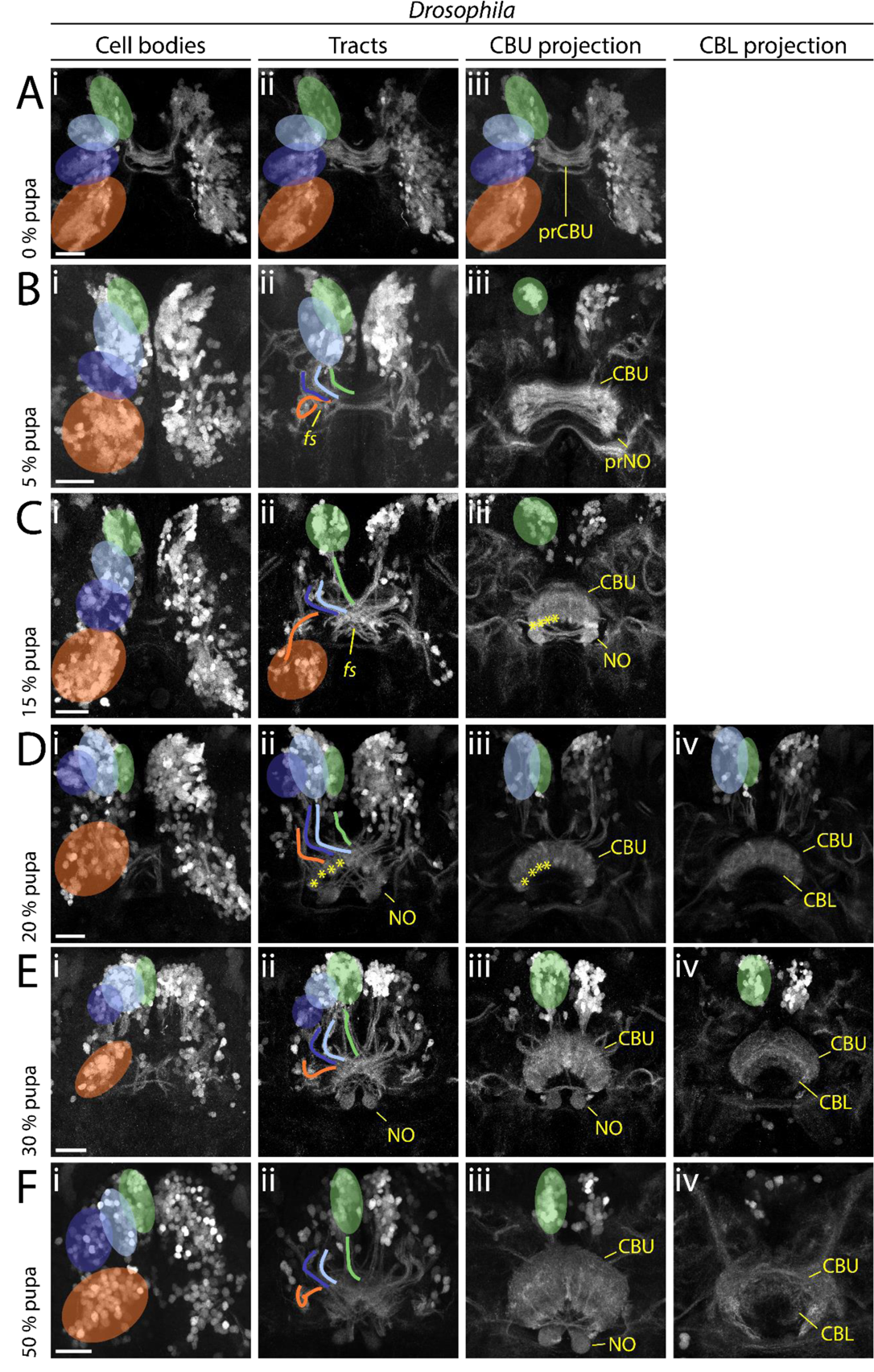
In *Drosophila*, the main developmental event of fascicle switching with resulting columnar fiber organisation occurs in the pupa. Displayed are sub-projections of an anti-GFP staining of the same brain per time point, to display the development and positioning of cell clusters (i) belonging to the DM1-4 lineage and their tracts (ii) (DM1 green, DM2 light blue, DM3 dark blue, DM4 orange) and final projections into the developing central complex neuropils (CBU iii, CBL iv). **(A-C)** Fascicle switching starts at 5 % and is very visible at 15 % pupation with the CBU and NO developing as result. **(D-F)** Fascicle switching continues, with the CBL developing. Also, the cell bodies get shifted, resembling the shape of the PB as result in later pupal stages. Following events are highlighted: Fascicle switching (*fs*) of DM1-3 was visible from 5 % onwards (B^ii^, C^ii^), with the formation of four columns of the CBU per hemisphere (asterisks in C^iii^ D^ii^, D^iii^). Abbreviations like in previous figures; fs fascicle switching event. Scale bars represent 25 µm.

First EGFP signal clearly corresponding to a CBU was found at 15 and 20 % (Fig. 8C/D^iii^) coinciding with the emergence of synapsin staining (Fig. 7F^ii^/H^ii^). We detected no pronounced projection into the CBL until 20 % while later projections remained low in intensity (Fig. 8F^IV^). Strong projections into the noduli were detectable from 15 % onwards (Fig. 8C^iii^). Following single tracts within the central complex was not possible.

In *Tribolium* pupal brains, the cell bodies of the Rx expressing DM1-4 groups remained comparably similar because they had undergone the respective re-arrangement already mostly in the embryo, and partially in the larva. From 0-15 % onwards, DM1-4 cells formed tracts, which underwent pronounced fascicle switching (Fig. 9A-C^ii^). The resulting division into columns became visible by the presence of strongly marked tracts from 0 % onwards in the CBU (Fig. 9A^iii^) and from 30 % in the CBL (Fig. 9E^iv^).

**Fig. 9:**
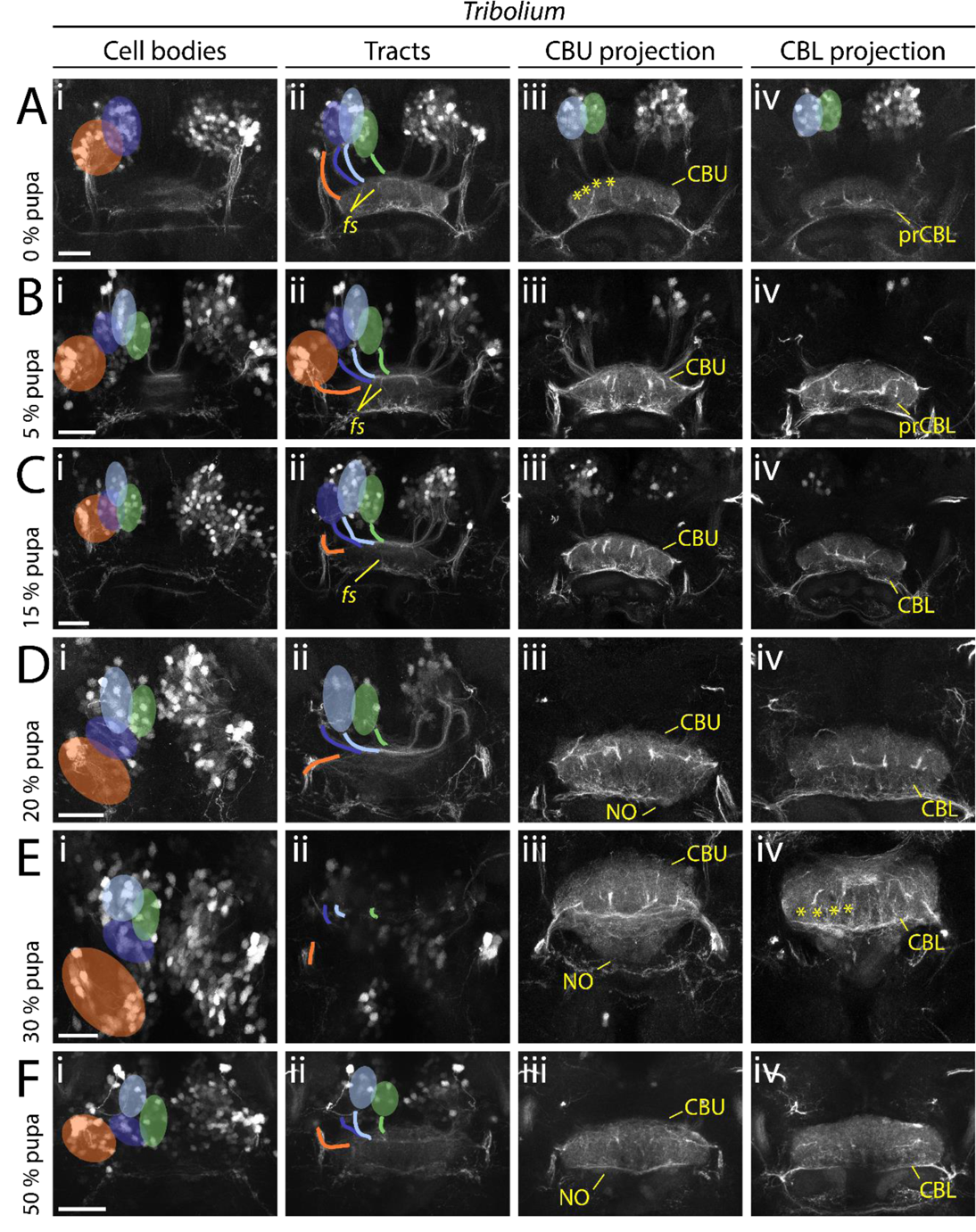
*Tribolium* pupal development illustrates how the adult central body becomes distinct from the larval form. Displayed are sub-projections of an anti-GFP staining of the same brain per time point, to display the development and positioning of cell clusters (i) belonging to the DM1-4 lineage and their tracts (ii) (DM1 green, DM2 light blue, DM3 dark blue, DM4 orange) and final projections into the developing central complex neuropils (CBU iii, CBL iv). **(A-C)** Fascicle switching becomes immediately prominently visible at 0 % and shows a columnar division of the CBU, and increases in later stages. **(D-F)** In later pupal stages, decussated projections go into the NO, and a column divided CBL. Following events are particularly highlighted: Fascicle switching (*fs*) of DM1-3 was visible from 0 % onwards (A^ii^, B^ii^, C^ii^), with a resulting formation of four columns of the CBU and CBL per hemisphere (earliest visible in A^iii^ and E^iv^, marked by asterisks). Abbreviations like in previous figures. Scale bars represent 25 µm.

Hence, we note an interesting pattern of decussation in the *Tribolium* L1 and pupal brains: While a decussated pattern was found in *Tribolium* L1 based on acetylated α-tubulin staining, it was not yet visible with the *Tc-rx-*EGFP line. In contrast, in the pupa, decussations of the *rx genetic neural lineages* became clearly visible in both species. It is therefore possible that the *rx genetic neural lineage* performs decussation postembryonically in both species, while other lineages perform this process already in the *Tribolium* embryo. We cannot exclude, however, that decussations of single neurites may not have been resolvable by the *Tc-rx-*EGFP line.

## Discussion

### Complex pattern of heterochronies and paedomorphocline of central complex development

Initially, the term *heterochrony* described differences in size and shape emerging mainly from different growth parameters such as rate and duration of growth (60, 64). *Sequence heterochrony* was introduced for cases where certain developmental steps change their position within a developmental sequence (65, 67). To assess the nature and complexity of central complex heterochrony, we used fifteen events of central complex differentiation for which we determined the absolute and relative timing in *Drosophila* and *Tribolium* development (Fig. 10), a two-dimensional approach of events and time, as previously proposed (67).

**Figure 10:**
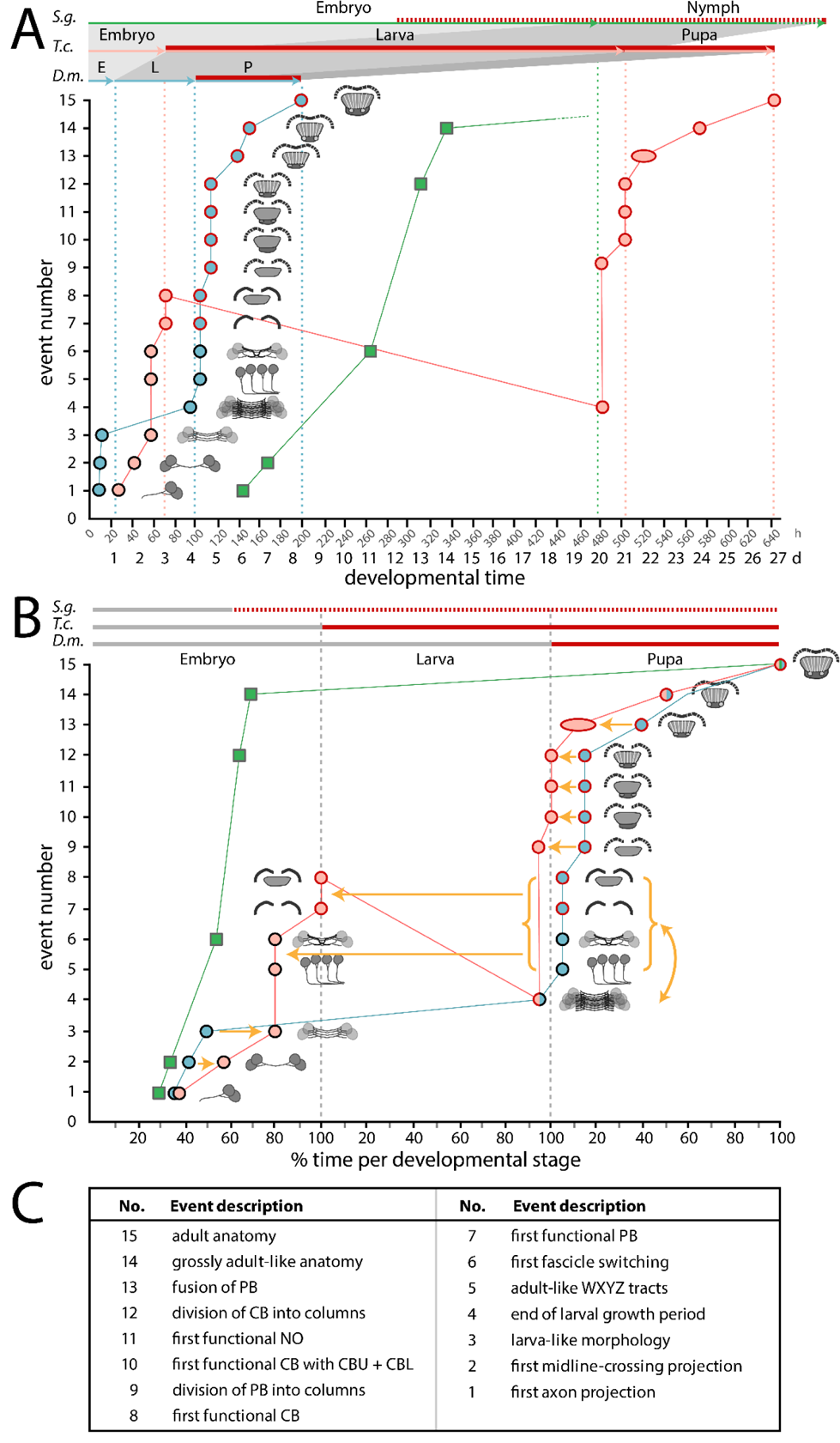
S**c**hematic summarizing the timing of developmental events of central complex heterochrony. Developmental time is depicted on the x axis as absolute time in hours and days (A) or relative time in % development of the respective life stages (B). Fifteen discrete events of central complex development (description in C and definition in Table S5) are depicted on the y axis and visualized with small sketches. The sequence of events reflects *Drosophila* development. The developmental trajectory shown for *Drosophila* (*D.m.*, blue) and *Tribolium* (*T.c.*, orange) is based on this work while *Schistocerca* (*S.g.*, green) is based on (81,114,122,123). Red contours of the circles and red lines on the top axes indicate presence of synapsin as a proxy for functionality of the central complex. Synapsin expression data was not available for *Schistocerca*, therefore neuromodulator expression was used instead (red hatched line). **(A)** A comparison on an absolute time scale highlights the large differences in actual time between species, and the resulting divergences over which period a respective animal has a functional central complex neuropil. *Drosophila* has the shortest generation time with the embryonic stage 33 %, the larval stage 17 % and the pupal stage being 71 % of the *Tribolium* time (32°C in *Tribolium*, 25°C in *Drosophila*). *Schistocerca* (∼31°C) embryonic central complex development takes more than double of the time of entire *Drosophila* central complex development (480 h versus 200 h). **(B)** Initial embryonic development leads to a heterochronic delay in *Tribolium* (orange arrows of events 2 and 3). In *Drosophila*, the larval growth phase follows (4) before in the pupa WXYZ tracts, decussation and gain of synapsin in PB and CB occur (5–8). Strikingly, these latter events are shifted into *Tribolium* embryogenesis. Further, we observed a sequence heterochrony, where the *Tribolium* larval growth phase occurs after events 5-8 instead of before like in *Drosophila* (curved yellow arrow and red line with negative slope). Pupal events 9 to 13 are heterochronically shifted to earlier stages of development in *Tribolium*. **(C)** Events are shortly described here and defined in Table S5.

We find a complex pattern of heterochronies, most of which reflect simple shifts in timing of differentiation events (orange arrows in Fig. 10). Interestingly though, some events occur earlier in *Drosophila* (e.g. first embryonic steps 1-3 – see Fig. 10 and Table S5) while with respect to others, *Tribolium* develops faster (steps 9 to 13). Importantly, some steps are even shifted between life stages: Formation of adult-like WXYZ tracts, first decussation and gain of functionality of the protocerebral bridge and central body are embryonic events in *Tribolium* but metamorphic events in *Drosophila* (steps 5-8).

We observe that ‘growth heterochrony’ (i.e. different timing, reduction or prolongation of growth, (64)) may not play a major role in central complex evolution because most of the growth happens at similar phases in both species (i.e. during early embryogenesis and during the larval stage). This contrasts with the crucial role that growth heterochrony was shown to play in the evolution of brains in other contexts. For instance, in humans, postnatal growth of the brain is strongly increased compared to chimpanzees (70). Across Mammalia an increase of proliferation rates probably led to gyrification (folding) of the cortex (117, 118). An intraspecific case of growth heterochrony has been noticed in insect castes, where bee queen brains develop faster and are larger as a result than worker bee brains (119).

Overall, the observed heterochronies reflect a paedomorphocline, i.e. an evolutionary juvenilization along the clades investigated (61). The *Schistocerca* central complex represents the ancestral situation while the *Tribolium* L1 central complex is paedomorphic as it shows similarity to a stage at 60% embryogenesis in *Schistocerca* where decussations have just initiated (81). Likewise, the *Drosophila* L1 central complex is paedomorphic to the *Schistocerca* neuropil, but its primordium equals an even earlier embryonic stage of about 45 to 50% (81), consisting of parallel fibers only.

### An example for sequence heterochrony in brain development

One of our key findings is the presence of sequence heterochrony that contributes to the different forms of larval central complex primordia in *Tribolium* versus *Drosophila*. Specifically, adult-like WXYZ tracts, fascicle switching and gain of functionality of protocerebral bridge and central body (steps 5-8) occur before main net growth of the central body in larvae of *Tribolium* while they occur after this larval growth period in *Drosophila*. To our knowledge, this is the first example of sequence heterochrony contributing to the evolution of brain diversity. Sequence heterochrony was previously described with respect to processes where sequences covered for example the entire development of crustaceans (120) or the different order of events of central nervous system, skeletal and muscular development in Metatheria and Eutheria (121).

The cell behavior underlying sequence heterochrony may be reflected in the development of *pointed-*positive DM1-4 cells in *Drosophila* (54, 55): During embryogenesis, their parallel midline-crossing neurites form the larval FB primordium where they arrest development. Only during late larval and early pupal stages, they continue development building decussations and projections into columns within the FB, forming pontine neurons. Hence, the homologous cells of *Tribolium* would just need to overcome the developmental arrest in order to form first decussations in the embryo. Imaging lines marking the *pointed* genetic neural lineage tailored by genome editing (28) would allow testing this hypothesis.

### An immature developmental stage of the central complex gained functionality in holometabolous larvae

It has been assumed that the larval central body of Tenebrionid beetles corresponds to the upper division (CBU, FB) of the adult. This assumption was based on its bar shape, the presence of tracts presumably prefiguring the lower division and some neuromodulator expression (20,33,51). We find that the functional central body of the L1 does not reflect any adult structure but is like a developmental stage that gained functionality (as defined by synapsin expression). It is immature with respect to the size and number of neurites of the neuropil, the pattern of decussation, the lack of columns and layers and in that the protocerebral bridge is not yet fused at the midline. Hence, *Tribolium* has two distinct forms of a functional central complex, one for the larval and one for the adult life stage. It should be noted that we cannot exclude that precursors of central complex neuropils could be present at earlier stages in form of synapsin-positive areas fused with other brain neuropils or as simple tracts that are not yet functional but present. However, given that central complex function is based on the intricate pattern of interconnections between central complex neuropils, we assume that a similar set of neuropils is required for a functional central complex.

How can a developmental stage become functional? Our data indicates that a connection of protocerebral bridge and central body by WXYZ tracts and some degree of decussation are required because all these events are specifically shifted into embryogenesis in *Tribolium*. This would be in line with current views on the function of the adult central complex where the intricate pattern of interconnections by columnar neurons, their decussations and resulting projections are required for integrative central complex functions like sky compass orientation (39,40,44,91). However, basic functionality does appear not to require the separation of upper and lower division of the central body nor a prominent columnar architecture. The question remains in how far the larval central complex function actually mimics the adult one. It is plausible that the simplified architecture leads to a less complex functionality.

### Evolutionary scenario – gain of a larval central complex in holometabolous larvae?

Hemimetabolan insects like *Schistocerca* reflect the ancestral situation where the entire central complex develops during embryogenesis. It was suggested that the situation in beetles and other holometabolous insect larvae (with their partial central complex) was less derived than the one of *Drosophila*, where a functional central complex emerges only during metamorphosis (33). However, our results reveal that *Tribolium* diverges from the other two in that its central complex gains functionality at an immature developmental stage.

This unexpected pattern may be explained by two different scenarios: First, a simple comparison between these three species would indicate that flies have retained the ancestral condition while beetle and other holometabolous insect larvae have gained functionality as evolutionary novelty. However, the larvae of beetles, which move and orient in an environment using eyes and legs are thought to be more similar to the ancestral holometabolous insect larva than the *Drosophila* maggot with its reduction of legs, eyes and head (124). Further, this scenario requires assuming independent gains of functionality of developmental stages in several holometabolous insect taxa. We prefer a second scenario, which puts the evolution of a functional larval central complex at the basis of the evolution of Holometabola. The holometabolan larva is an immature but functional life stage. This is reflected by a number of immature organs like simplified legs, antennae and eyes while other organs lack completely (e.g. wings) (124). Minimal functionality of the central complex might have been required for the evolution of the larval stage for guiding at least some basic behavior involving eyes and legs. In this scenario, the distribution of larval functional central complexes in several taxa would reflect conservation while the lack in *Drosophila* larva would reflect a loss as evolutionary divergence. Indeed, the fly maggot may need less elaborate orientation behavior because it hatches within the food source which usually supports its entire development. Unfortunately, data on embryonic central complex development is missing for most taxa calling for respective studies in order to test our scenario (20,51,125,126).

### Does the larval central complex recycle phylogeny?

Similarity has been noted between the larval central complex of *Tenebrio molitor* and the larval structure in the Branchiopod *Triops cancriformis* with its rudimentary protocerebral bridge and non-columnar central body (12, 127). Moreover, there is a striking similarity of the *Tribolium* larval central complex as we describe it with the adult crayfish central complex, with its bilateral WXYZ tracts projecting into a uniform non-layered, non-columnar central body, and its V-shaped, only slightly fused protocerebral bridge (128, 129), as well as with some shrimp central complexes (131). Hence, the holometabolan larval central complex could reflect a phylogenetic intermediate that occurred in the evolution towards the insect central complex. However, in that case, ontogeny does not simply reflect phylogeny. Rather, the functional larval central complex of holometabolous larvae represents a regression to an evolutionary precursor. 100 years after Haeckel’s death one might be inclined to state ‘ontogeny reflects – or recycles – phylogeny’ and heterochrony may be a driving force.

## Material and Methods

### General considerations

We adhered to the nomenclature presented in (82), except for our reference to the DM4 ipsilateral fascicle as tract, which we remain with the term W tract (75). In addition, we referred to central body divisions as upper and lower division, instead of fan-shaped and ellipsoid, because these terms were used in classical literature and the lower division has an ellipsoid shape only in a few species. Similarly, we use the tradtional term ‘columns’ for vertical subdivisions in the central complex while ‘slices’ has been suggested as synonym (82).

Animals were kept at 32°C for *Tribolium castaneum* and 25°C for *Drosophila melanogaster* under respective standard conditions (132, 133). Execpt for embryos and young larvae where sexing was not possible, females were selected for stainings. Besides in Fig. 5G (N=1), Fig. 5H (N=2) and Fig. 6B-D (N=2), the dataset consisted of at least N=3 tissues. All stacks from which figures were created and films in .avi format thereof can be found under figshare (https://figshare.com/account/home#/projects/64799). All *Drosophila* and *Tribolium* stocks, antibodies and dyes, as well as primers are documented in Tables S2-S4. Detailed information on all methods used can be found in the Supporting Material and Methods.

### Tc-Rx antibody generation and verification

The anti-*Drosophila* Rx antibody was kindly gifted by Dr. Uwe Walldorf (98). No cross reactivity to the Tc-Rx protein was found. Hence, we generated an antibody against Tc-Rx by cloning the region N-terminal to the homeobox domain into a GoldenGate vector containing a SUMO peptide (KNE001, Supporting Material and Methods), expressing it in BL21-DE3 Rosetta bacteria and purifying it by immobilized metal ion affinity chromatography. A guinea pig antibody was then raised against the purified peptide by Eurogentec (Kaneka Eurogentec S.A., Belgium). Finally, specificity of the antibody was verified by in situ hybridisation against *rx* RNA combined with Tc-Rx immunostaining as well as immunostaining of *Tc*-*rx* RNAi-mediated knockdown embryos (Fig. S1).

### *rx*-EGFP transgenic lines

For *Drosophila*, a trangenic line marking large parts of *rx* expression was not available. Therefore, we generated a bicistronic line by CRISPR/Cas9 mediated homology-directed repair (Fig. S3)(28). Towards this end, we removed the endogenous STOP codon of the *rx* ORF to generate an in-frame *rx*-*EGFP* fusion gene. In addition, we included a sequence encoding for the P2A peptide between the *rx* ORF and *EGFP* CDS to ensure that two distinct proteins from a common RNA will have been translated (for information on the P2A peptide see (108, 109)). We also included an eye marker allowing us to screen G_1_ positives with ease. The repair template was cloned using the Gibson assembly kit (New England Biolabs, MA, USA). Suitable target sites without off-targets were identified using the CRISPR Optimal Target Finder (134) (http://targetfinder.flycrispr.neuro.brown.edu/). Respective guides were cloned into an U6:3-*Bbs*I vector and subsequently tested by a T7 Endonuclease I assay. The repair template and guideRNA containing plasmids were co-injected into *Act5C-Cas9, DNAlig4[169]* embryos (135). Surviving G_0_ animals were crossed individually to *w^-^* virgins of the opposite sex and the G_1_ generation was screened for eye marker and EGFP reporter. The overlap of EGFP and Rx was determined by double immunostainings in adults and embryos. Indeed, we found that each cell expressing Rx now also expressed EGFP, largely located in the cytoplasm.

For *Tribolium*, we identified a suitable transgenic line in the GEKU base website where its insertion had been mapped to the upstream region of *Tc*-*rx* (# E01101, http://www.geku-base.uni-goettingen.de/, Fig. S2)(136). This *Tc-rx*-EGFP line was verified by Rx/GFP co-immunostainings which revealed that all EGFP expressing cells also expressed Rx (with the exception of the eye transformation marker).

Both, *Dm-rx*-EGFP and *Tc*-*rx*-EGFP, were made homozygous and all data used derives from homozygous stocks.

### Comparative staging and determining central complex events

A description of the stages that we defined are documented in the Supporting Material and Methods and Table S5. Exact values for the timing of central complex developmental events displayed in Fig. 10 are found in Table S5.

### Fixation, staining, imaging and image processing

Fixation, in situ hybridization and immunostainings were performed as described in (27, 32) with details in the Supporting Information. Images were taken with a Leica SP8 confocal microscope (Wetzlar, Germany) with standard settings. Images were examined using Fiji software (137). 3D reconstructions were performed using Amira 5.4.1 (Visage Imaging, Fürth, Germany) and figures created using Adobe Illustrator CS5 (Adobe Systems, San José, CA, USA).

## Author contributions

GB and MSF designed experiments, analyzed the data and wrote the paper. MSF conducted most experiments. GB conceived of the project idea. KNE and MSF designed and conducted experiments with respect to antibody production and transgenic line generation. All authors approved the final version of the manuscript.

## Acknowledgements

Dr. Felix Quade helped with 3D reconstructions and Lara Markus provided some embryonic and larval immunostainings. We thank Prof. Uwe Walldorf for providing the Dm-Rx antibody, and Prof. Christian Wegener for providing the anti-synapsin antibody. Dr. Achim Dickmanns supported protein expression and purification. Analyses of brain anatomy and homologous cell group identification were supported by Prof. Volker Hartenstein. We want to further thank Dr. Stephen H. Montgomery and Prof. Robert A. Barton for fruitful discussions. We thank Drs. Marita Buescher and Nico Posnien for valuable discussions as well as Stefan Dippel for sharing unpublished data on pupal staging, Elke Küster and Claudia Hinners for technical support and Dr. Elisa Buchberger for helpful corrections of the manuscript. MSF and KNE were supported by the Göttingen Graduate Center for Molecular Biosciences, Neurosciences and Biophysics (GGNB).

