## Supporting Information for "Sequence heterochrony led to a gain of functionality in an immature stage of the central complex: a fly-beetle insight"

### Supporting Figures and Tables

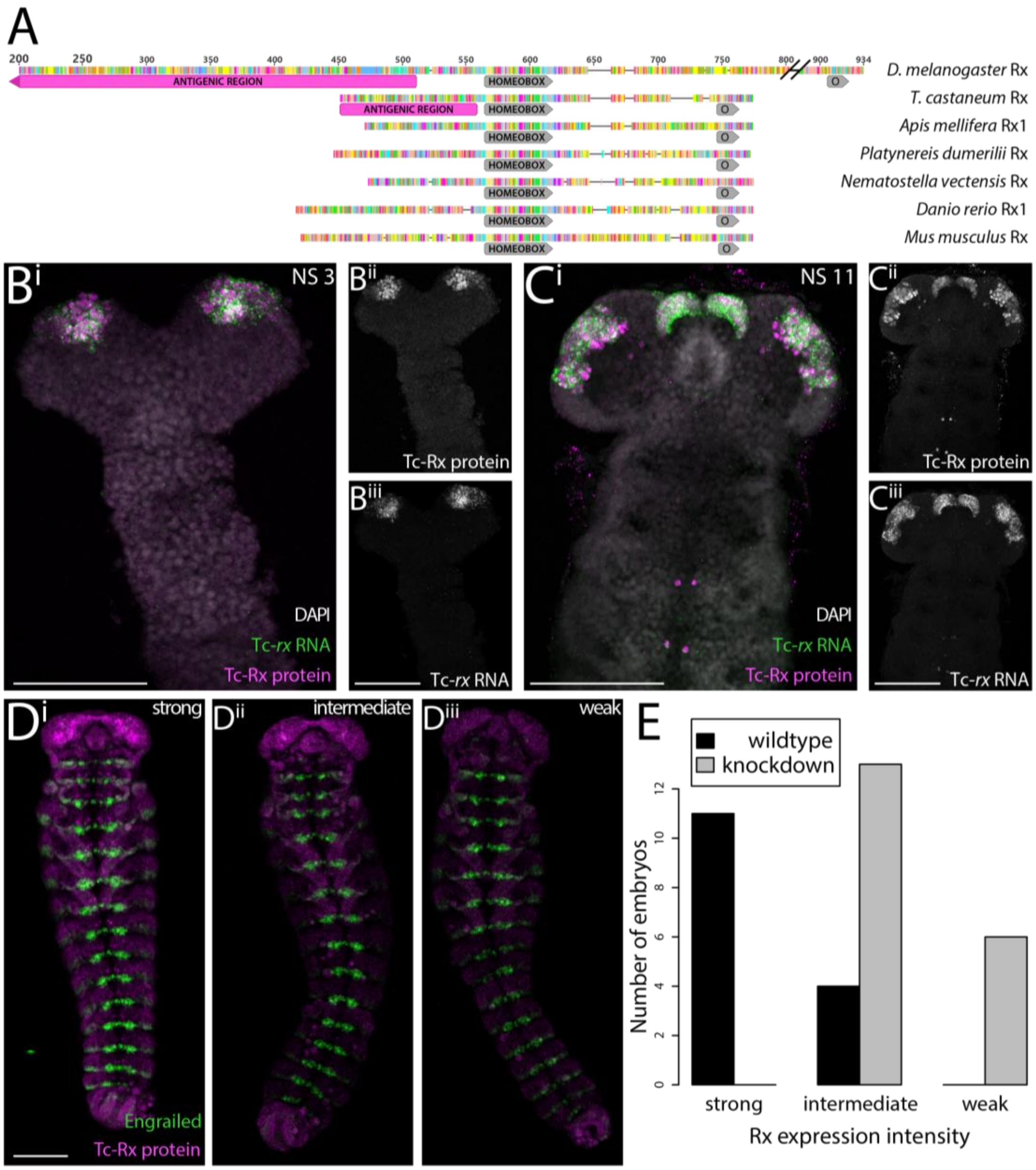


**S1 Figure: Generation and validation of the Tc-Rx antibody. (A)** Alignment (Geneious 11.1.5, Geneious Alignment) of Rx proteins of *Drosophila* and *Tribolium* as well as representative species. The conserved homeobox and OAR (O) domains (grey) are present in all proteins. Antigenic regions for the Dm-Rx (1,2) and the Tc-Rx antibody are displayed in magenta. The Dm-Rx protein was shortened for better display (amino acids 1 to 200 and most between 800 and 900 are not displayed). Notice that the *Drosophila melanogaster* (*D. melanogaster*) antigenic region appears to be absent in *Tribolium castaneum* (*T. castaneum*) and all other species. **(B-C)** Tc-Rx protein and *Tc-rx*

RNA expression in *Tribolium* embryos of neurogenesis stages 3 and 11 (3) were depicted (Zeiss LSM510, 40x immersion objective) as maximum intensity projections (DAPI for structure as average projection). Anterior is to the top. Animals were mounted dorsal up. The signal detected in the antibody staining against Tc-Rx protein (magenta) overlapped to a high degree with the signal detected in the in situ hybridization (green). Note that while the protein of Tc-Rx was located in the nucleus, *Tc-rx* RNA was also in the cytoplasm of the cell soma which resulted in a different cellular localization. **(D)** To validate the specificity of the Tc-Rx antibody, we performed a RNAi mediated *Tc-rx* knockdown. Indeed, Tc-Rx expression was reduced in knockdown embryos. Depicted are three categories of Tc-Rx expression (i.e. Tc-Rx antibody staining intensity, magenta, as maximum intensity projections) after knockdown (strong, equaling wildtype, in $D^{i}$, intermediate in $D^{ii}$, weak in $D^{iii}$). To accommodate for differences in intensity of staining, a co-staining against Invected/Engrailed with the respective antibody was performed. **(E)** 34 RNAi embryos were categorized into the three expression intensity groups in a blinded experiment. Wildtype animals showed a high level of expression and were mostly grouped in category ‘strong’ with some in category ‘intermediate’. No knockdown animals were grouped into the ‘strong’ category, most in ‘intermediate’ and some in ‘weak’ (Fisher‘s exact test, P<0.001). Scale bars represent 100 µm.

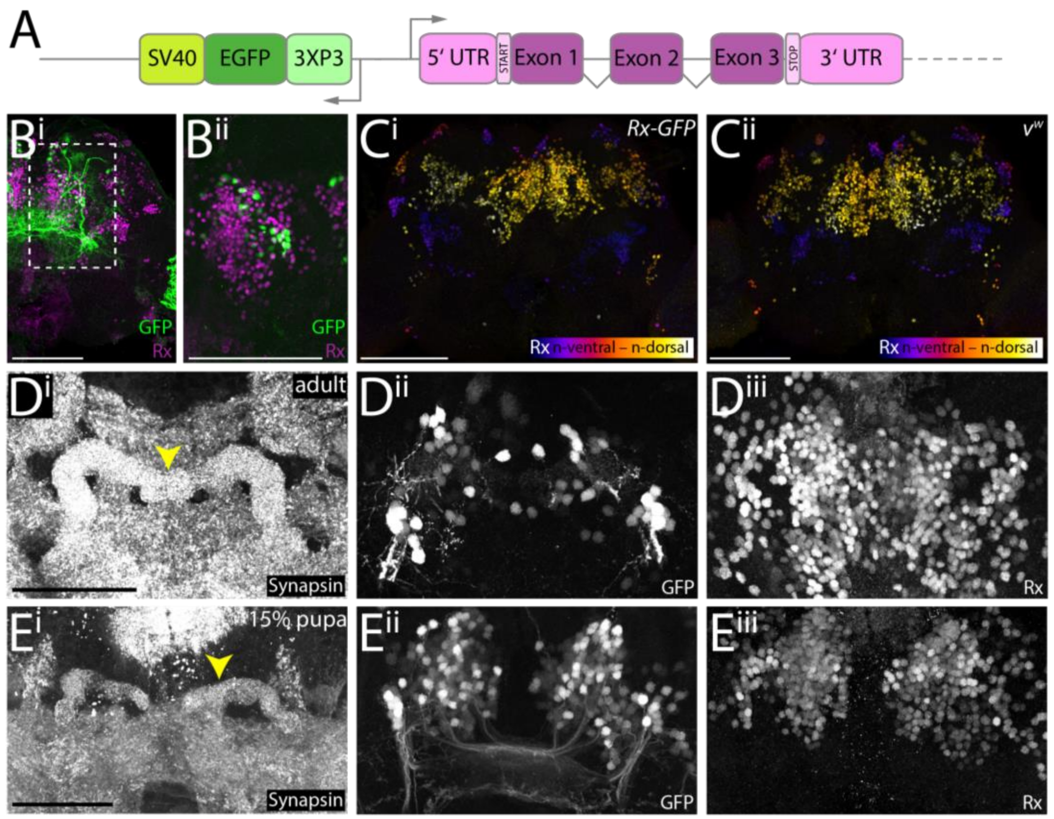


**S2 Figure: Characterization and validation of *Tribolium rx-EGFP* enhancer trap line. (A)** The *Tribolium rx-EGFP* enhancer trap was taken from the GEKU screen collection (4) where enhancer traps were generated by *piggyBac*-mediated transposition. A *3XP3-EGFP-SV40* cassette randomly inserted upstream of the *Tc-rx* gene in opposite direction (insertion site mapped by (4)). **(B)** Maximum intensity projections of immunostainings against GFP and Tc-Rx in adult brains of the *Tc-rx*-EGFP line. The line only marked a small subset (approximately 5-10 %) of Tc-Rx expressing cells in the adult. This also applies to the n-dorsal region ($B^{ii}$). However, all GFP expressing cells also expressed Tc-Rx. Coexpression was verified manually. **(C)** The introduction of the enhancer trap cassette did not visually influence Tc-Rx expression, as domains were highly similar between transgenic Rx-GFP ($B^{i}$) and wildtype *vermillion-white* ($v^{w}$, $B^{ii}$, (5)) animals, as visualized by color-coded maximum intensity projections. Observed qualitative differences in Tc-Rx expression in the transgenic or wildtype condition (N=3 each) were approximately as large as the differences between the genetic backgrounds. **(D)** A crop of a maximum intensity projection of cells surrounding the adult protocerebral bridge (yellow arrowhead, $D^{i}$) shows the coexpression of GFP ($D^{ii}$) and Tc-Rx ($D^{iii}$) in a subset of cells that were subsequently used in this study. **(E)** An analogous analysis in young pupal brains of cells surrounding the protocerebral bridge ($E^{i}$) revealed more GFP expressing cells ($E^{ii}$) with overlap to Tc-Rx cells ($E^{iii}$) than in the adult (D). Scale bars in B and C represent 100 µm and in D and E 50 µm.

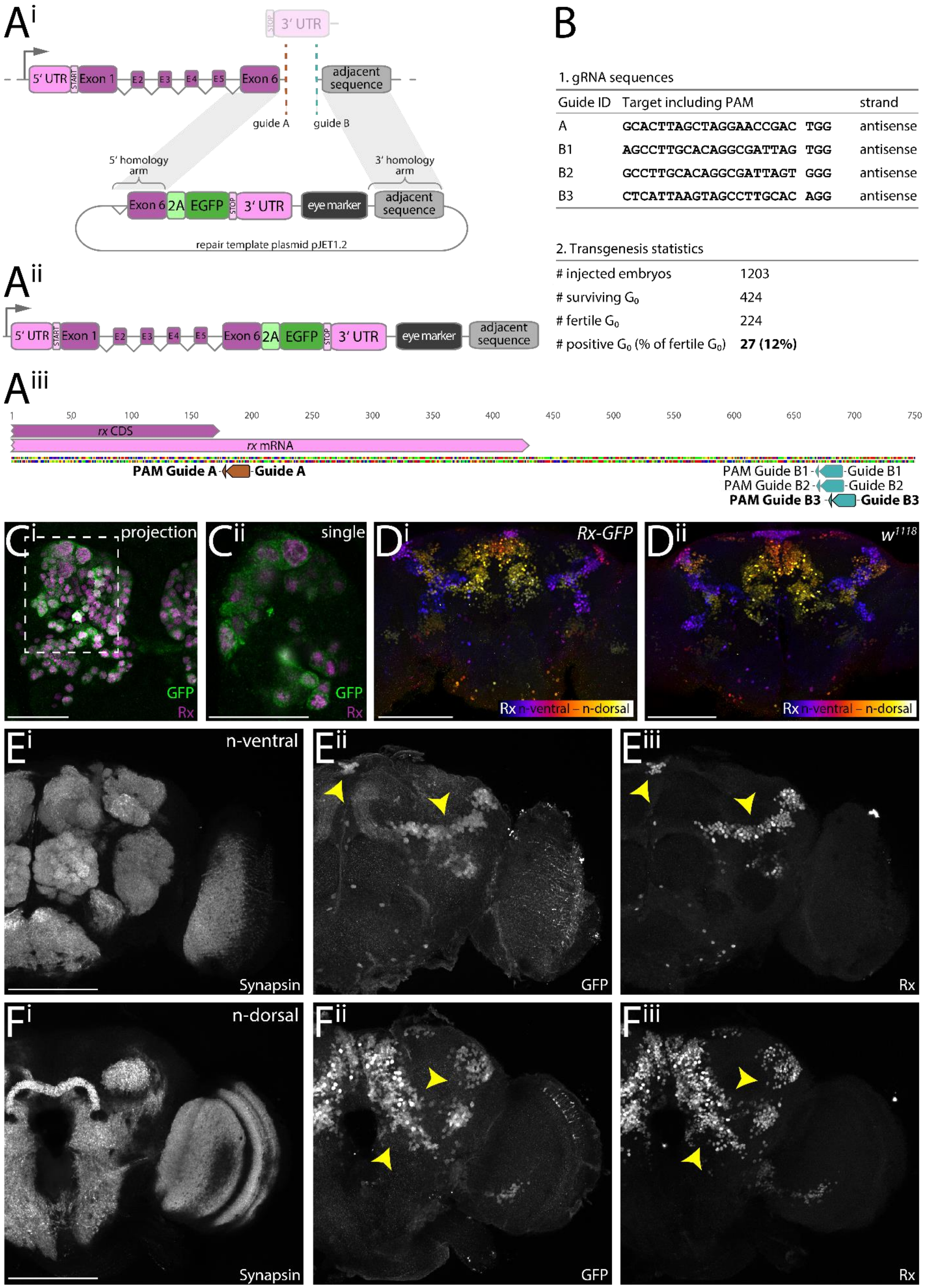


1. gRNA sequences

| Guide ID | Target including PAM | | strand |
|---|---|---|---|
| A | GCACTTAGCTAGGAACCGAC | TGG | antisense |
| B1 | AGCCTTGCACAGGCGATTAG | TGG | antisense |
| B2 | GCCTTGCACAGGCGATTAGT | GGG | antisense |
| B3 | CTCATTAAGTAGCCTTGCAC | AGG | antisense |

2. Transgenesis statistics

| | |
|---|---|
| # injected embryos | 1203 |
| # surviving $G_0$ | 424 |
| # fertile $G_0$ | 224 |
| # positive $G_0$ (% of fertile $G_0$) | **27 (12%)** |

**S3 Figure: Strategy, generation and validation of *Drosophila* bicistronic *rx*-EGFP transgenic line. (A$^{i}$)** Strategy of building a *Dm-rx*-EGFP line (modified from (6)). Two gRNAs next to the endogenous STOP codon (guide A, brown dashed line) and downstream of the *Dm-rx* 3’UTR (guide B, blue dashed line) were used. The DNA repair template included a sequence encoding for a *P2A* self-cleaving peptide, *EGFP*, the endogenous region between guide A and B (*Dm-rx* 3’UTR and a fraction of intergenic region), and the *3xP3-DsRed-SV40* eye marker, as well as 1 kb homology arms flanking the insertion sites. **(A$^{ii}$)** The edited transgenic locus comprises a common open reading frame of both *Dm-rx* and *EFGP* with a STOP after *EGFP*. **(A$^{iii}$)** Four gRNAs were used in different combinations to generate similar transgenic lines. The gRNAs used for the transgenic line used in this study are marked in bold (guide A and B3). **(B)** Overview of gRNA sequences and transgenesis statistics upon injection (7) for the *Drosophila* Rx-GFP transgenic line. **(C$^{i}$)** Immunostaining of anti-Dm-Rx (magenta) and anti-GFP (green) in the *Dm-rx*-EGFP line showed that all visible cells that expressed Dm-Rx also expressed GFP, shown in a smooth manifold extension (SME) projection (8) of a brain hemisphere of a S16 embryo. The region marked with a dotted line in C$^{i}$ is shown in **(C$^{ii}$)** as a single slice. Here, the different cellular localizations are visible. Dm-Rx retained its nuclear localization, while GFP located to the cytoplasm, demonstrating functionality of the P2A peptide. **(D)** The transgenic line had normal Dm-Rx expression, shown by anti-Dm-Rx immunostaining and depth color-coded maximum intensity projection in the Rx-GFP line (D$^{i}$) and the origin wildtype strain $w^{1118}$ (D$^{ii}$). Observed qualitative differences in Dm-Rx expression in the transgenic or wildtype condition (N=3 each) were approximately as large as the differences between the genetic backgrounds. **(E-F)** Dm-Rx and EGFP expression matched in adult brains (see yellow arrowheads for exemplary double-positive areas). Maximum intensity projections of synapsin immunostainings (E$^{i}$, F$^{i}$), GFP (E$^{ii}$, F$^{ii}$) and Dm-Rx (E$^{iii}$, F$^{iii}$) in an adult *Drosophila* brain. Anti-synapsin (E$^{i}$, F$^{i}$) marked brain position. E is n (neuraxis)-ventral and F is n-dorsal (9). Scale bars in D-F represent 100 µm and in C 25 µm.

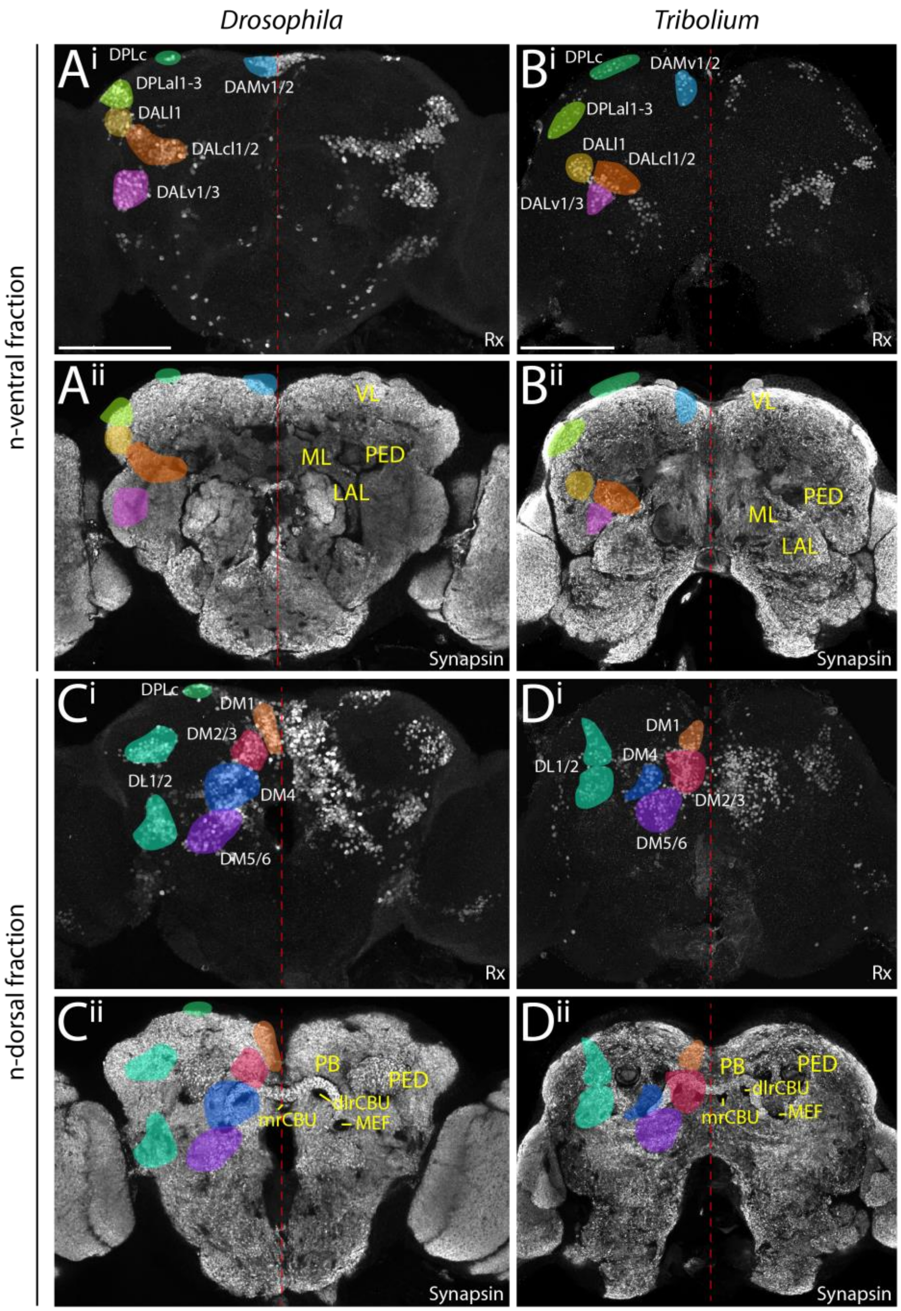


**S4 Figure: Conserved expression of Rx protein in the adult brain of *Drosophila melanogaster* (A, C) and *Tribolium castaneum* (B, D) as well as lineages marked by Rx expression.** We mapped the labeled Rx-positive cells to previously described lineages of the *Drosophila* brain using locations relative to other brain structures and their projection pattern as criterion ((10,11); www.mcdb.ucla.edu/Research/Hartenstein/dbla/index.html and references therein). We tentatively named *Tribolium* cell clusters by using similar locations and projections as compared to the *Drosophila* atlas, used as guide. A list of all lineages with names and descriptions can be found in Table S1.

Hemispheres are separated by a red dotted line for orientation. Due to the cell body rind expression of Rx, domains and proposed lineages could be separated into two fractions, n-ventral and n-dorsal, corresponding to each half of the insects' brains. For each species, one image stack was used and separated into two fractions. Rx expression is displayed by a maximum intensity projection of a sub-stack of an anti-Rx immunostaining (i). Basic anatomical structure of the insects' brains is displayed by a SME projection (8) of a synapsin immunostaining (ii). On this projection, in the left hemisphere the locations of the proposed lineages are shown color-coded, while on the right hemispheres, basic anatomical structures are annotated that assist understanding differences in domain position between the species (yellow). Abbreviations: VL vertical lobe, ML medial lobe, PED peduncle, LAL lateral accessory lobes, mrCBU medial root of the upper division of the central body, dlrFB dorso-lateral root of the CBU, PB protocerebral bridge, MEF medial equatorial fascicle. Scale bars represent 100 µm.

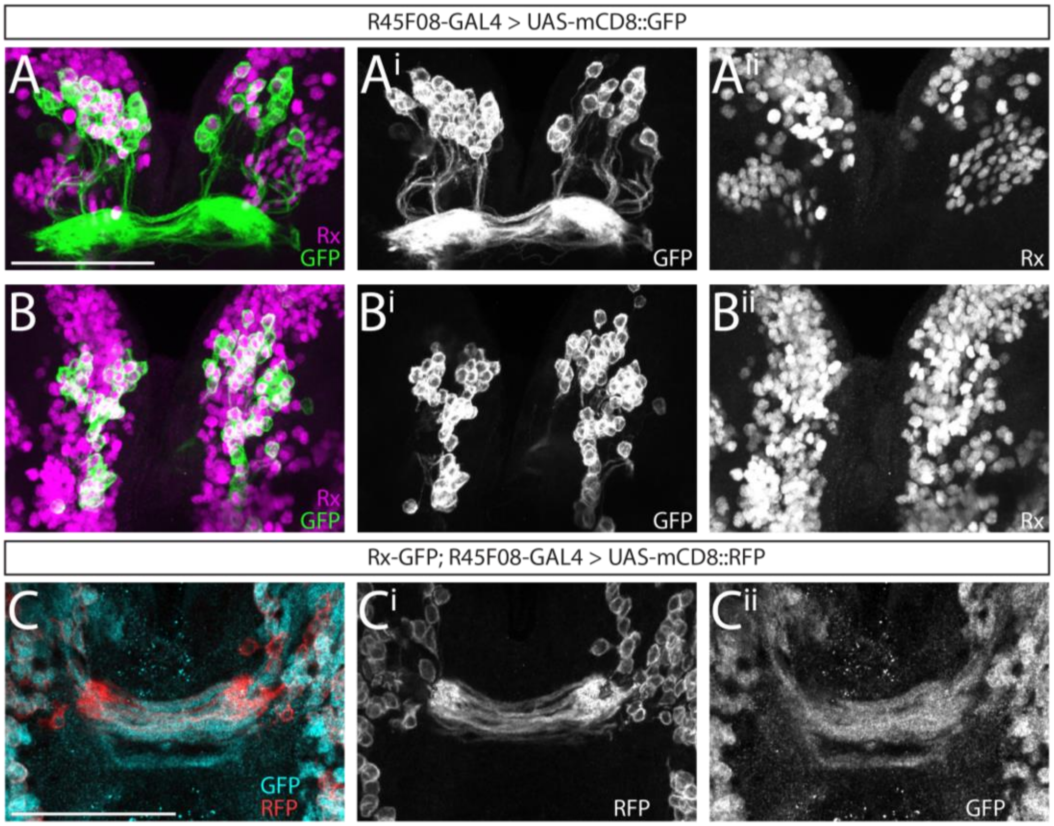


**S5 Figure: Previously described *pointed*-positive cells of the central complex are a subset of Dm-Rx-positive cells.** Displayed is a co-localization of Dm-Rx-positive neural cells and cells under the control of R45F08-GAL4 (12,13) shown in brains of *Drosophila* wandering third instar larvae. **(A-B)** Antibody staining in a cross of the R45F08-GAL4 line and UAS-mCD8::GFP was performed against Dm-Rx (depicted in magenta) and GFP (green) to reveal the coexpression of cell bodies of lineages DM1-3/6, marked through the R45F08-GAL4 line, and Dm-Rx. Approximately 90% of the R45F08-GAL4 GFP positive cells were Dm-Rx-positive as well (A-A$^{ii}$ first half, B-B$^{ii}$ second half of the stack). **(C)** Antibody staining in animals (N=2) of the respective cross from subcrosses of the Rx-GFP line each with R45F08-GAL4 line and the UAS-mCD8::RFP (SMEs, see (8)). This resulted in a coexpression of GFP in a Dm-Rx expression pattern and RFP under control of R45F08-GAL4. Antibody staining against GFP (cyan) and RFP (red) revealed coexpression of both fluorescent proteins in midline crossing projections. Although RFP is membrane-bound and GFP cytoplasmic, there were several fascicles showing coexpression of RFP and GFP. This corroborated the high degree of overlap of Dm-Rx and DM1-3/6 lineage offspring shown in panels A and B. Scale bars represent 50 µm.

**S1 Table: Proposed lineages expressing Rx in the adult *Drosophila* (*Dm*) and *Tribolium* (*Tc*) brain.** Listed are eleven lineages with identifier, name and a description relative to the neuroaxis, as well as the position in Fig. 2 and S4 Figure and the degree how unequivocally the assignment of their stereotypical projections was. Identification of lineages is based on (10,11), https://www.mcdb.ucla.edu/Research/Hartenstein/dbla/index.html, and references therein. Abbreviations: PED peduncle, LAL lateral accessory lobes, AVLP anterior ventrolateral protocerebrum, SLP superior lateral protocerebrum, SMP superior medial protocerebrum, PB protocerebral bridge, MEF medial equatorial fascicle, CA calyx.

| Lineage identifier (alternative) | Lineage name | Description (relative to neuraxis) | Fraction | *Dm:* projections identifiable? | *Tc:* projections identifiable? |
|---|---|---|---|---|---|
| DALcl1/2 | dorso-anterior lateral, centro-lateral 1/2 | n-ventro-lateral to PED, n-anterior to LAL | n-ventral | yes | no |
| DALl1/2 | dorso-anterior lateral, lateral 1/2 | n-anterior to AVLP, n-ventral and lateral to PED | n-ventral | yes, DALl1 | no |
| DALv1/3 | dorso-anterior lateral, ventral | n-ventral to AVLP, lateral to LAL | n-ventral | no | no |
| DPLal1-3 | dorso-posterior lateral, antero-lateral 1-3 | lateral to anterior SLP | n-ventral | partially, DPLal2/3 | no |
| DPLc | dorso-posterior lateral, central | n-anterior to posterior SLP | n-ventral | yes | no |
| DAMv1/2 | dorso-anterior medial ventral 1/2 | n-anterior to SMP | n-ventral | yes | no |
| **DM1 (DPMm1)** | **dorso-medial 1 (dorso-posterior medial, medial 1)** | n-anterio-dorsal (Tc)/n-anterio-ventral (Dm) and medial to PB | **n-dorsal** | **yes** | **yes** |
| **DM2/3 (DPMpm1/2)** | **dorso-medial 2/3 (dorso-posterior medial, postero-medial 1/2)** | n-dorso-medial to PB | **n-dorsal** | **yes** | **yes** |
| **DM4 (CM4)** | **dorso-medial 4 (centromedial 4)** | n-posterio-lateral to PB, n-dorsal to MEF | **n-dorsal** | **yes** | **yes** |
| DM5/6 (CM1/3) | dorso-medial 5/6 (centromedial 1/3) | n-posterio-lateral to PB, n-posterio-dorsal to MEF | n-dorsal | no | no |
| CP2/3 (DL1/2) | centroposterior 2/3 (dorsolateral 1/2) | n-posterio-lateral to CA and n-dorsal to posterior PED | n-dorsal | yes | yes |

**S2 Table: Primer list.** P1 to P12: black writing – annealing part, red – overlapping part, green – PAM modification.

| Name | Sequence | Purpose |
|---|---|---|
| Tc-rx-N_fw | ATGGAATCGGACCGTTGTGAAG | protein expression |
| Tc-rx-N_rev | CTTGCATCCGTCTCCCTC | protein expression |
| Golden Gate linker sequence fw | CCAGGTCTCATGGT | protein expression |
| Golden Gate linker sequence rev | GGGGGTCTCCTCGAGTCA | protein expression |
| GG_ccdB_F | ACATGATTGCGGCGTTGCC | KNE001 vector |
| GG_ccdB_R | TGTCTCTCGAGGAGACCGTCGACCTGCAGACT | KNE001 vector |
| GEKU-Rx-GFP_wt_fw | AGTTGCGAGATGTGCGAGT | homozygous stock generation |
| GEKU-Rx-GFP_wt_rev | CGTCCAGACTTGCCACTTTG | homozygous stock generation |
| GEKU-Rx-GFP_trans_rev | CTCTAAAATAAGGCGAAAGGC | homozygous stock generation |
| Tc-rx-probe-fw | ATGGAATCGGACCGTTGTGAAGA | full length rx probe |
| Tc-rx-probe-rev | GCAGTCCTTTGGTGATGTTCTCC | full length rx probe |
| I_P1_Back-F1 | CCGGATGGCTCGAGTTTTTCAGCAAGATCACATCGCCTGGGATGCG | rx bicistronic construct |
| I_P2_Rev_F1 | GACAATGGATACCATTCCCTTGTTCAGG | rx bicistronic construct |
| I_P3_F1-F2 | CCTGAACAAGGGAATGGTATCCATTGTCGGGTCCGGCGCCACCAAC | rx bicistronic construct |
| I_P4_Rev_F2-F3 | GTGAACAGCTCCTCGCCCTTGCTCACCATGGGGCCGGGGTTCTCCTCC | rx bicistronic construct |
| I_P5_F2-F3 | GACGTGGAGGAGAACCCCGGCCCCATGGTGAGCAAGGGCGAGGAG | rx bicistronic construct |
| I_P6_Rev_Back-F3 | AGAATATTGTAGGAGATCTTCTAGAAAGATCTACTTGTACAGCTCGTCCATGCCGAG | rx bicistronic construct |
| II_P7_Back-F4 | CCGGATGGCTCGAGTTTTTCAGCAAGATCGTTAGTCGGTTCCTAGCTAAGTG | rx bicistronic construct, PAM of guide A modified |
| II_P8_Rev_F4-F5 | CTCTAATTGAATTAGATCACATACGATTAGTATAACAGATAAGCATTCC | rx bicistronic construct, PAM of guides B1-3 modified |
| II_P9_F4-F5 | GCTTATCTGTTATACTAATCGTATGTGATCTAATTCAATTAGAGACTAATTCAATTAGAGC | rx bicistronic construct, PAM of guides B1-3 modified |
| II_P10_Rev_F5-F6 | CATTAAGTAGCCTTGGATACATTGATGAGTTTGGACAAAC | rx bicistronic construct |
| II_P11_F5-F6 | GTCCAAACTCATCAATGTATCCAAGGCTACTTAATGAGTTGATTAATAAG | rx bicistronic construct |
| II_P12_Rev_Ba-F6 | GAATATTGTAGGAGATCTTCTAGAAAGATGTTCTTTCAATTTGTAAGACATAGGTTTTTAG | rx bicistronic construct |
| III_P13_Rev_F3 | CTACTTGTACAGCTCGTCCATGCCGAG | rx bicistronic construct |
| III_P14_F3-F4 | CTCGGCATGGACGAGCTGTACAAGTAGCGTTAGTCGGTTCCTAGCTAAGTG | rx bicistronic construct |
| DmRx_CDS_3'UTR_fw | CGTCTCTGCCACTAATTAGACAGC | rx SNP sequencing |
| DmRx_CDS_3'UTR_rev | GAATAGACTTCTTCGTCAGCCG | rx SNP sequencing |
| DmRx_3'UTR_int-region_fw | CGTGTTGTAAGTACATATTTCTGAGGCAG | rx SNP sequencing |
| DmRx_3'UTR_int-region_rev | CTTGAGGAGCGAGGCACAC | rx SNP sequencing |
| DmRx_trans-ver_fw | GTCGCCGCAGAACCTGAG | rx molecular screening |
| DmRx_trans-ver_rev | CATGGAGCCAGTAGTTCATGC | rx molecular screening |
| DmRx_trans-ver_nested_fw | CATAGAACTGCTCGATGTGG | rx molecular screening |
| DmRx_trans-ver_nested_rev | GATTCAACTGCGGCTACTGC | rx molecular screening |
| DmRx_trans_seq_Ct_fw | GACTGGCAAGGGTTCGAG | rx molecular screening |
| DmRx_trans_seq_iRe_rev | CATGTGAGTCCTTTGTTTGC | rx molecular screening |

116 **S3 Table: List of primary and secondary antibodies as well as dyes used in this study.**

| Antibody name | Antigen / Immunogen | Origin species | Source | Dilution |
|---|---|---|---|---|
| anti-Dm-Rx | *Drosophila* Rx N-terminal fragment | rabbit | gift from Dr. Uwe Walldorf (Saarbrücken, Germany), Davis et al. 2003 | 1:1000 |
| anti-Tc-Rx | *Tribolium* Rx N-terminal fragment | guinea pig | this paper | 1:750 |
| anti-Engrailed 4D9 | Engrailed/invected (Immunogen: Invected (C-terminal two-thirds of the invected protein); recombinant) | mouse | gift from Dr. Marita Buescher (Göttingen, Germany), DSHB | 1:10 |
| chk-anti-GFP | GFP (*Aequorea victoria*) | chicken | ab13970, Abcam (Cambridge, UK), used in Supplementary Figure 3 and 6 | 1:1500 |
| rab-anti-GFP | GFP (*Aequorea victoria*) | rabbit | A11122, ThermoFisher Scientific/Invitrogen (MA, USA) | 1:1000 |
| anti-RFP | RFP (full length) | rabbit | ab62341, Abcam (Cambridge, UK) | 1:1000 |
| anti-Synapsin | Synapsin (Immunogen: GST-Synapsin-GST fusion protein expressed in E. coli and purified by glutathione affinity) | mouse | gift from Dr. Christian Wegener (Würzburg, Germany), DSHB | 1:15-1:40 |
| anti-α-acetylated Tubulin | α-acetylated Tubulin (Immunogen: acetylated tubulin from the outer arm of Strongylocentrotus purpuratus (sea urchin) | mouse | T7451, MERCK/Sigma-Aldrich (Darmstadt, Germany) | 1:40 |
| anti-rab-Alexafluor 488 | rabbit (Gamma Immunoglobins Heavy and Light chains) | goat | A11070, ThermoFisher Scientific/Invitrogen (MA, USA) | 1:1000 embryos, 1:500 other stages |
| anti-chk-Alexafluor 488 | chicken (Gamma Immunoglobins Heavy and Light chains) | goat | A11039, ThermoFisher Scientific/Invitrogen (MA, USA) | 1:1000 embryos, 1:500 other stages |
| anti-mou-Alexafluor 555 | mouse (Gamma Immunoglobins Heavy and Light chains) | goat | A21425, ThermoFisher Scientific/Invitrogen (MA, USA) | 1:1000 embryos, 1:500 other stages |
| anti-rab-Alexafluor 555 | rabbit (Gamma Immunoglobins Heavy and Light chains) | goat | A21430, ThermoFisher Scientific/Invitrogen (MA, USA) | 1:1000 embryos, 1:500 other stages |
| anti-rab-Alexafluor 647 | rabbit (Gamma Immunoglobins Heavy and Light chains) | goat | A21245, ThermoFisher Scientific/Invitrogen (MA, USA) | 1:1000 embryos, 1:500 other stages |
| anti-gp-Alexafluor 647 | guinea pig (Gamma Immunoglobins Heavy and Light chains) | goat | A21450, ThermoFisher Scientific/Invitrogen (MA, USA) | 1:1000 embryos, 1:500 other stages |

| Dye name | Source | Dilution |
|---|---|---|
| DAPI | D1306, ThermoFisher Scientific (MA, USA) | 1:1000 |

117

**S4 Table: *Drosophila* and *Tribolium* stocks used in this study.**

| Species | Stock name | Stock Number | Source | Description |
|---|---|---|---|---|
| *Drosophila* | **Rx-GFP** | **-** | **this paper** | **Genotype: $w^{1118}$, 3XP3-dsRED, $rx^{GFP}$; founder 374.2; primary line** |
| | $w^{1118}$ | e.g. 3605 | e.g. Bloomington Drosophila Stock Center | used for control experiments (S3 Figure) |
| | Act5C-Cas9, Lig4[169] | 58492 | Bloomington Drosophila Stock Center | (7), Cas9 line used for generation of Rx-GFP bicistronic line (S3 Figure) |
| | $w^{1118}$; $wg^{Gla\text{-}1}$/CyO | - | Wimmer Department, Göttingen (gift) | used to generate homozygous stocks of the Rx-GFP bicistronic line (S3 Figure) |
| | R45F08-Gal4 | 49565 | Bloomington Drosophila Stock Center | (12,13); used to determine overlap of lineages DM1-3/6 to Rx expressing cells (S5 Figure) |
| | 20xUAS-mCD8::GFP | 32194 | Bloomington Drosophila Stock Center | Chromosome 3, used to determine overlap of lineages DM1-3/6 to Rx expressing cells (S5 Figure) |
| | UAS-mCD8::ChRFP | 27392 | Bloomington Drosophila Stock Center | Chromosome 3, used to determine overlap of lineages DM1-3/6 to Rx expressing cells (S5 Figure) |
| *Tribolium* | **Rx-GFP** | **E01101** | **GEKU-base** | **ref (4), primary line** |
| | $v^{w}$ | - | ref (5) | $vermilion^{white}$, (5), used for control experiments (S2 Figure) |

**S5 Table: Description and definition of fifteen central complex related events used in this study to illustrate heterochronic development in *Tribolium castaneum* (*Tc*), *Drosophila melanogaster* (*Dm*) and *Schistocerca gregaria* (*Sg*).** Events were defined by using our dataset of anti-GFP and anti-synapsin stainings with both species, to determine potential differences between them, and by using the central complex literature as reference point. Time points for each event are included, as absolute time in hours and relative time per developmental period in percent. Note that length of embryonic developmental periods was taken from (3,14) and Scholten and Klingler (unpublished), stages were determined using morphological criteria and then time points were calculated from these works. Larval and pupal developmental times were determined specifically for our *rx* transgenic lines and pupal time points were then verified by morphological criteria using works by Dippel (unpublished) and (15). Information on *Schistocerca* central complex events as well as relative and absolute developmental time was taken from (16–19). Abbreviations: PB protocerebral bridge, CB central body, CBU upper division of the central body, CBL lower division of the central body, NO noduli, *Tc Tribolium castaneum*, *Dm Drosophila melanogaster*, *Sg Schistocerca gregaria*.

| Event No. | Short description | Definition | Time point in *Dm* | Time point in *Tc* | Time point in *Sg* |
|---|---|---|---|---|---|
| **1** | first axon projection | The first axonal projection of Rx-positive cells in the prospective central complex region | 8.8h/37% | 28h/39% | 144h/30% |
| **2** | first midline-crossing projection | The first projection of Rx-positive cells that spans across the midline | 10.3h/43% | 42h/58% | 168h/35% |
| **3** | larva-like morphology | A pattern of projections and cell body location that allows clear identification of DM1-4 lineage origin and is thus similar to the larval pattern | 12.2h/51% | 58h/81% | |
| **4** | end of larval growth period | The end of larval growth at the end of development of the last larval instar, correlated with an increase in size of central complex structures but without notable changes in morphology | 95.3h/95% | 482.4h/95% | |
| **5** | adult-like WXYZ tracts | The presence of identifiable WXYZ tracts, i.e. axonal fiber bundles corresponding to DM1-4, that project in a similar morphological way and shape as in the adult, i.e. first n-posterior, then towards the midline in a region corresponding to a central body neuropil | 104h/5% | 58h/81% | |
| **6** | first fascicle switching | The first occurrence of fascicle switching, causing a decussated fiber pattern, i.e. X-shaped crossings in the central complex | 104h/5% | 58h/81% | 264h/55% |
| **7** | first functional PB | The first synapsin-positive structure identifiable as protocerebral bridge | 104h/5% | 72h/100% | |
| **8** | first functional CB | The first synapsin-positive structure identifiable as central body | 104h/5% | 72h/100% | |
| **9** | division of PB into columns | Presence of vertical subdivisions in the PB, apparent in an anti-synapsin staining | 114h/15% | 482.4h/95% | |
| **10** | first functional CB with CBU + CBL | The first synapsin-positive structure identifiable as a lower division of the central body (or ellipsoid body), i.e. the first division into lower and upper central body part | 114h/15% | 504h/0% | |
| **11** | first functional NO | The first synapsin-positive structure identifiable as noduli | 114h/15% | 504h/0% | |
| **12** | division of CB into columns | Presence of vertical subdivisions in the CB region primarily based on patterns of the *rx* transgenic line (anti-GFP) and secondarily a heterogenous distribution of synapses, most importantly the absence of synapsin in otherwise anti-GFP positive regions (anti-synapsin) | 114h/15% | 504h/0% | 312h/65% |
| **13** | fusion of PB | The fusion of the protocerebral bridge at the midline | 139h/40% | 511-532h/5-20% mean: 521.5h/12.5% | |
| **14** | grossly adult-like anatomy | An anatomy that grossly resembles the adult pattern, particularly with respect to the DM1-4 cell bodies and tracts | 149h/50% | 574h/50% | 336h/70% |
| **15** | adult anatomy | Mature central complex anatomy of the adult | 199h/100% | 644h/100% | 1848h/100% |

**S6 Table: Stages and their definition included in this study.**

| *Drosophila* | | | *Tribolium* | |
|---|---|---|---|---|
| **Stage** | **Time (h)** | **Description** | **Time (h)** | **Description** |
| **Embryos** | variable | All staging follows (14) | variable | Staging to 48 h: (3); >48 h: Scholten and Klingler (unpublished) |
| **L1** | ≤ 1 after hatching | selection after removing any previously hatched larvae on apple agar plate | ≤ 1 after hatching | selection by falling through 300 µm gaze sieve on which embryos were kept |
| **50 % (mid-) larva** | 37.5 | timing started after selection like for L1, end of L2 larval development | ~ 216 (9 d) | timing started after selection like for L1, approximately L4 stage |
| **95 % (late-) larva** | 70-75 | up to event 2 (15), crawled to top, no movement | 410-432 (17-18 d) | last larval stage, stemmata migration started, see Ho 1961 |
| **0 % pupa** | 0 | up to event 8 (15), particularly shortened body, white puparium | 0 | stemmata migration ended (medial position near vertex), see (40) |
| **5 % pupa** | 5 | up to event 14 (15), operculum ridge visible, abdominal gas bubble | 7 | 2 rows of ommatidia, shiny cuticle, see (40), and Dippel (unpublished) |
| **15 % pupa** | 15 | up to event 25 (15), particularly anterior bubble, expelled armature | 21 | 4 rows of ommatidia, mandible tip sclerotized, see (40), and Dippel (unpublished) |
| **20 % pupa** | 20 | up to event 26 (15), criteria of 15 % and prominent Malphigian tubules | 28 | 4-6 rows of ommatidia, mandible tip sclerotized, see (40), and Dippel (unpublished) |
| **30 % pupa** | 30 | up to event 27 (15), criteria of 20 %, yellow body, eyes not included | 42 | 6 rows of ommatidia, mandible tip sclerotized, see (40), and Dippel (unpublished) |
| **50 % pupa** | 50 | criteria of 30 %, otherwise only time | 70 | 7 rows of ommatidia, see (40), and Dippel (unpublished) |
| **adult** | ≤ 12 after eclosion | eclosed with signs of virgin females (light body coloring) | ≤ 24 after eclosion | eclosed with light brown body coloring |

**S7 Table: Immunohistochemistry in stages (excluding embryos) of both species.** There are two variations of adult stainings. Antibodies were used as in Table S3 except for synapsin. PB phosphate buffer (20), T Triton-X-100 with % in PB, PFA paraformaldehyde, NGS normal goat serum.

| Preparation steps | L1 larvae | Larvae | Pupae | Adults |
|---|---|---|---|---|
| **Fixation** | 1 h on ice, in 4 % PFA | 50 % larva: 1 h on ice, in 4 % PFA, other: 1.5 h on ice, in 4 % PFA | 1.5 h on ice, in 4 % PFA | 1.5-2 h on ice, in 4 % PFA |
| **Post-fixation washes** | 1 rinse, 3 30 min washes in PB-T 0.1 % | 50 % larva: 1 rinse, 3-4 30 min washes in PB-T 0.1 %, other: 1 rinse, 3-4 30 min washes in PB-T 0.3 % | 1 rinse, 3-4 30 min washes in PB-T 0.2-0.3 % | I: 1 rinse, 3-4 30 min washes in PB-T 0.3 %, II: 1 rinse, 3-4 30 min washes in PB-T 0.5 % |
| **Blocking** | o/n at 4°C in 4 % NGS in PB-T 0.1 % | o/n at 4°C in 5 % NGS in PB-T 0.1 % (50% larva), or 0.3 % (other) | o/n at 4°C in 5 % NGS in PB-T 0.2-0.3 % | I: o/n at 4°C in 5 % NGS in PB-T 0.5 % II: 24 h at 4°C in 5 % NGS in PB-T 0.3 % |
| **First antibody** | 4 h at RT in 2 % NGS in PB-T 0.1 %, Synapsin 1:30 (Dm), 1:20 (Tc) | 4-6 h at RT in 2 % NGS in PB-T 0.1 % (50%), 0.3 % (other), Dm Synapsin 1:30, Tc 1:20-30 | 5 h at RT or 40-48 h at 4°C in 2 % NGS in PB-T 0.2-0.3 %, Synapsin 1:25 (Dm), 1:15 (Tc) | I: 6 h at RT in 2 % NGS in PB-T 0.5 %, II: 72 h at 4°C in 2 % NGS in PB-T 0.3 %, Synapsin 1:25 (Dm), 1:15 (Tc) |
| **Post-1st antibody washes** | 1 rinse, 3 30 min washes in PB-T 0.1 % | 1 rinse, 4 30 min washes in PB-T 0.1 % (50%), 0.3 % (other) | 1 rinse, 4 40 min washes in PB-T 0.2-0.3 % | 1 rinse, 4 50 min washes in PB-T 0.3/0.5 % |
| **Secondary antibody** | o/n at 4°C in 2 % NGS in PB-T 0.1 % | o/n at 4°C in 2 % NGS in PB-T 0.1 % (50%), 0.3 % (other) | o/n to 24 h at 4°C in 2 % NGS in PB-T 0.2-0.3 % | I: 24 h at 4°C in 2 % NGS in PB-T 0.5 % II: 48 h at 4°C in 2 % NGS in PB-T 0.3 % |
| **Post-2nd antibody washes** | 1 rinse, 1 30 min wash including DAPI, 1 rinse, 3 30 min washes, all in PB-T 0.1 % | 1 rinse, 1 30 min wash including DAPI, 1 rinse, 3 30 min washes, all in PB-T 0.1 % (50%), 0.3 % (other), 2 h wash in PB | 1 rinse, 1 30 min wash including DAPI, 1 rinse, 3 30 min washes, all in PB-T 0.2-0.3 %, 2 h wash in PB | 1 rinse, 1 30 min wash including DAPI, 1 rinse, 4 30 min washes, all in PB-T 0.3/0.5 %, 3 h wash in PB |
| **Embedding medium** | VectaShield H-1000 (Vector Laboratories) | RapiClear 1.47 (SUNjin Lab, Hsinchu City, Taiwan) | RapiClear 1.47 (SUNjin Lab, Hsinchu City, Taiwan) | RapiClear 1.47 (SUNjin Lab, Hsinchu City, Taiwan) |

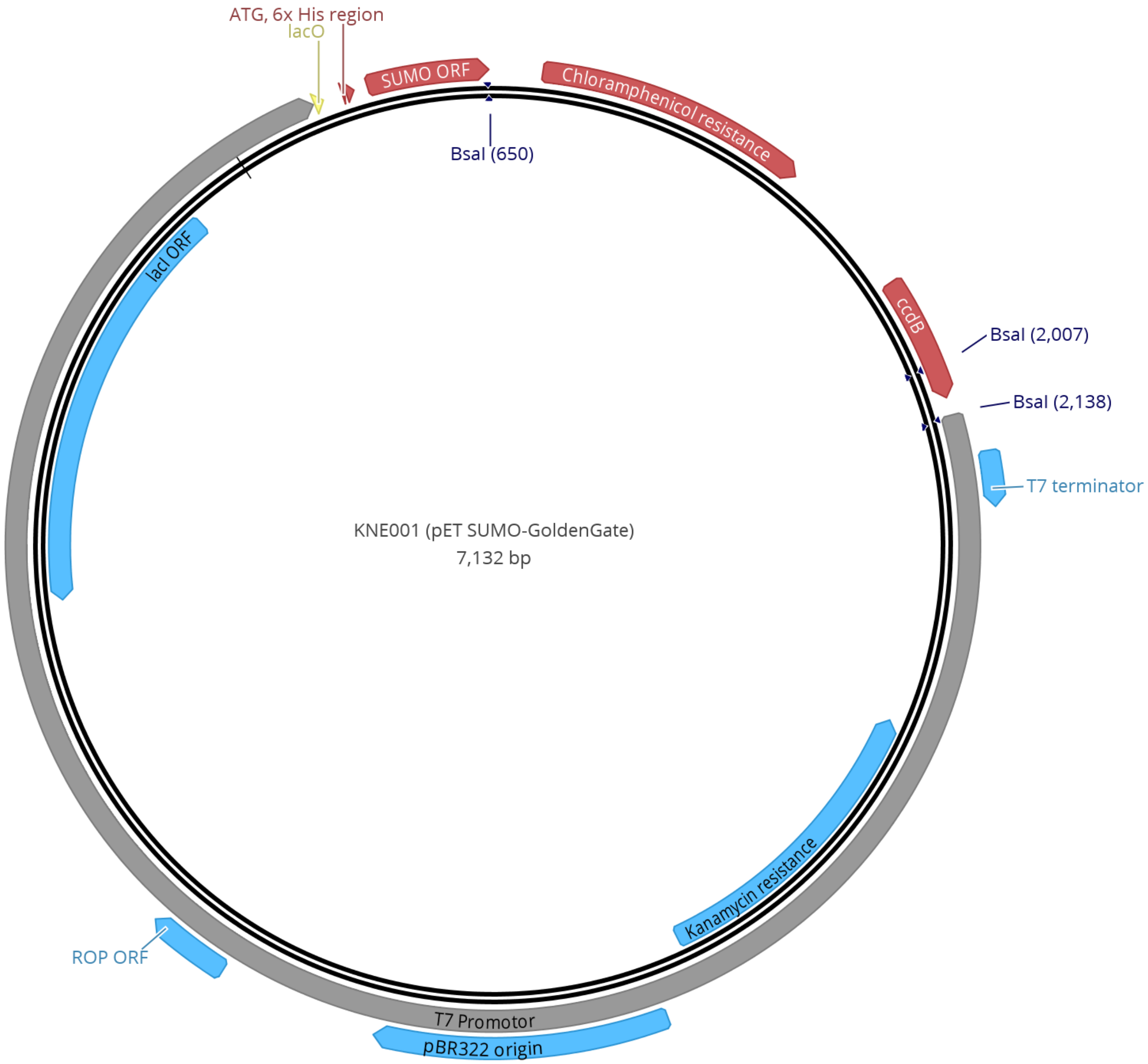


**Vector map S1: KNE001 vector map** (displayed with Geneious 11.1.5, https://www.geneious.com).

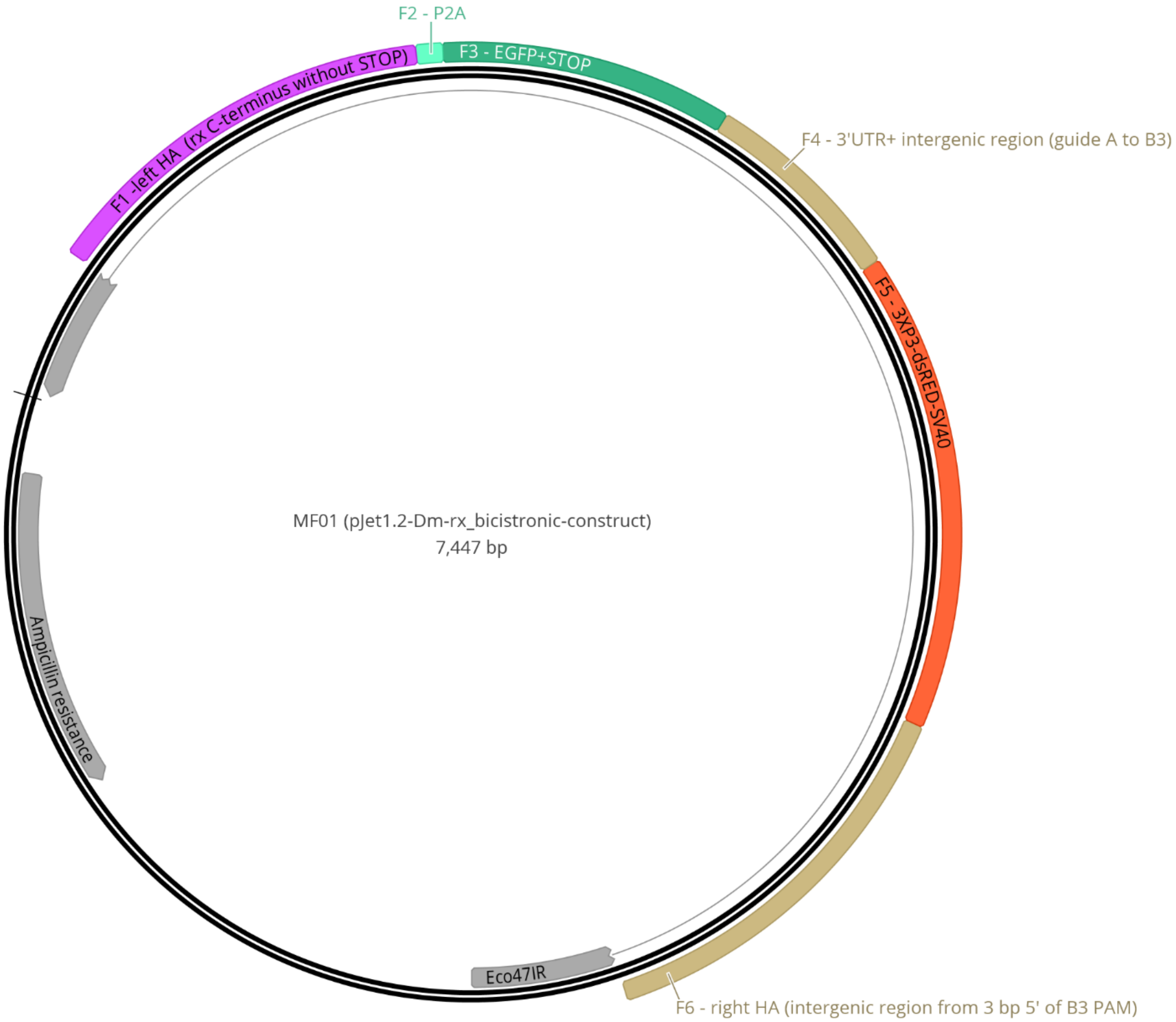


**Vector map S2: MF01 vector map** (displayed with Geneious 11.1.5, https://www.geneious.com). In the construct, we included an insect codon-optimized version of the P2A peptide (21), with following sequence: GGGTCCGGCGCCACCAACTTCTCCCTGCTGAAGCAGGCCGGCGACGTGGAGGAGAACCCCGGCCCC.

### Supporting Results

**Mapping of Rx-positive cell groups to known lineages of the insect adult brain**

We aimed at determining to which previously described lineages the Rx-positive cells belonged. The lineages in *Drosophila* had been described as the published atlas (10,11). We reassigned them in the *Drosophila* brain and transferred *Drosophila* knowledge to the *Tribolium* brain. Assignments of conserved Rx expressing cell groups in the cell body rind in both species' brains were based on two aspects. First, synapsin staining revealed common synapse-rich neuropils as well as synapse-absent tracts and fascicles that can be homologized between the two species. With this, domains of the *Tribolium* brain could be linked to domains and known lineages in *Drosophila*. Second, since Rx-positive lineages were defined by stereotypical projections, an additional antibody staining against GFP in the characterized Rx transgenic lines (S2 and S3 Figures) revealed some lineage-typical projections. Therefore, projections helped in some cases to verify lineage identity beyond cell body position. However, for most lineages, projections were not distinguishable. We identified eleven lineages in both species that cover most of the Rx expressing cell groups in the adult brain (DALcl1/2, DALl1/2, DALv1/3, DPLal1-3, DPLc, DAMv1/2, DM1 (DPMm1), DM2/3 (DPMpm1/2), DM4 (CM4), DM5/6 (CM1/3), CP2/3 (DL1/2, S4 Figure, S1 Table): In the *Tribolium* brain, all n-ventral lineages were not marked by projections through our transgenic line. They have been identified due to the basic anatomical position of the cell bodies that was very similar to Rx expressing lineages in the *Drosophila* brain. In the *Drosophila* Rx-GFP line all n-ventral lineages were – if at all – only faintly marked by projections. Visible were projections of the lineage group DALc1/2 that projected n-posterior to the peduncle into the central complex, the likely dorsal tract of the DPLal2/3 lineage that projected into the superior lateral protocerebrum, the short projection of the DPLc1 sublineage and the dorso-medial projection of the DAMv1/2 lineage into the superior medial protocerebrum. In the n-dorsal fraction, both projections of hemilineages of the CP2/3 lineage were visible in both species, one reaching n-anterior over the peduncle and projecting into the superior medial protocerebrum, one starting n-posterior of the peduncle and projecting n-ventro-anterior to it. With the available tools, we could not determine homology of these lineages further. To verify this tentative lineage identification, based mostly on cell body location, specific transgenic lines need to be generated and subsequent antibody stainings need to be performed, particularly in *Tribolium*, to further reveal the characteristic projection patterns of each lineage.

**Description of Rx-positive subgroups of DM1-4 lineages in *Tribolium* and *Drosophila***

In addition to the general descriptions of cell body location and projections on a lineage level (Fig. 2-3, S4 Figure), DM1-4 lineages were previously divided into sub-groups and single tracts (11). We wanted to describe which of those sub-groups and tracts are visible in the *Drosophila* adult brain and describe analogous sub-groups and tracts in *Tribolium*. These groups were differently marked in the imaging lines in both species due to the different design of the transgenic lines (see S2 and S3 Figures). Note that the projections of individual tracts or neurons in the respective neuropils were hard to distinguish because a high number of cells were marked.

In *Drosophila*, the DM4 Rx expressing cell group consisted of three subgroups, one localized n-anterior, and two n-posterior to the lateral tip of the PB. They projected axons to form a common projection as part of the MEF which bifurcated near the midline, where parts went into a n-ventral midline crossing projection n-ventral to the whole CX. This projection might be partially shared by the upper intermediate tract of CM3 or the dorsal tract of CM1 (11). The other part projected mainly into the CBU (‘intermediate tract; (11). The DM3 Rx expressing group consisted of two groups, one more n-anterior, one n-posterior to the lateral side of the PB. The group’s axons formed parts of the dlrCBU together with DM2 in the ’anterior-ventral tract’ (11). Parts of these cells’ axons projected into the n-dorsal plexus (also CBUppl, or FBppl, see e.g. (13)), while substantial parts went in a more n-ventro-posterior part together with DM4. DM2 consisted of three groups, two n-anterior (one of which is more n-dorsal), one n-posterior to the PB. They projected together into the n-dorsal plexus of CBU (’anterior-ventral tract'; (11), slightly more n-dorsal than DM3. The projection bifurcated, one more n-anterior, one more n-posterior. The DM1 group consisted of three subgroups, all n-anterior to the protocerebral bridge. One more n-ventral and slightly more lateral, two were more n-dorsal, of which one was n-anterior to the other. The n-ventral group formed a separate more lateral projection (potentially the 'anterior descending tract'; (11)) in comparison to the common projection of the other group ('anterior-ventral tract'; (11)).

In *Tribolium*, DM4 consisted of two groups localized n-anterior to the PB tip and one n-posterior to the PB tip. The bifurcation of the tract from both groups was similar to *Drosophila*, and they thus could also share a projection with a CM3 tract. A division into an n-anterior and n-posterior part was similar to *Drosophila*. A third group present in *Drosophila* was not marked or was not present in *Tribolium*. DM3 consisted of two main groups, one more n-anterior to the PB, one n-posterior to the PB, an arrangement similar to *Drosophila*. They projected together with DM2 and 1 into the n-dorsal fraction of the CBU, while sharing the dlrCBU tunnel with DM3, with projections very similar to *Drosophila*. Cell bodies of DM2 were difficult to visualize but were slightly more medial to the DM3 belonging group. Hence approximate position was similar, but a subdivision in groups was hardly possible. Cell bodies of DM1 were sparse, with some n-anterior and some n-posterior to the protocerebral bridge, like *Drosophila* without a subdivision into groups possible. The projection into the CBU was very similar. Note that, in general, cell groups of DM1-3 were n-dorsal, and not like in *Drosophila* n-anterior to the PB.

### Supporting Material and Methods

**Tc-Rx antibody generation and verification**

An antibody for the *Drosophila* Rx (Dm-Rx) protein was kindly gifted by Dr. Uwe Walldorf (1). Its specificity was verified by absence of staining in Dm-Rx null mutant brains and by a similar expression pattern as *Dm-rx* RNA (1,2).

We tested cross-specificity of this antibody to the *Tribolium* Rx (Tc-Rx) protein. However, no staining was detected (data not shown). As the antigenic region of Dm-Rx used for antibody generation by (1) is absent or highly diverged in the *Tribolium* Rx protein (like in a number of other species, see Fig. S1) we used the Tc-Rx N-terminal region (amino acids 1-107), avoiding highly conserved homeobox and OAR domains to generate a suitable antibody. An N-terminal 321 bp gene sequence was amplified (primers including linker sequences: *Tc-rx-N_fw* and *Tc-rx-N_rev*, Table S2) from wildtype cDNA and cloned into a Golden Gate vector containing a 6x His-Tag and a sequence encoding for a SUMO polypeptide (KNE001, pET SUMO-GoldenGate) with a molar ratio of 1:5 of insert to vector (see below for source, modifications and cloning information).

For subsequent protein expression and purification, we essentially followed (22). The vector was transformed into bacteria of the BL21-DE3 Rosetta strain. Bacteria expressed the peptide in TB (Terrific Broth) medium with the addition of 15 mM Glucose by 0.8 mM IPTG induction at an $OD_{600}$ of 0.8 for four hours, were then harvested (5,000×g, 20 min, 4°C), resuspended in lysis buffer (50 mM Tris-HCl pH=7.5, 150 mM NaCl, 10 mM Imidazole), fractionated using a microfluidizer 110S (Microfluidics, MA, USA) and cell debris was removed by centrifugation (30,000×g, 30 min, 4°C). The peptide was subsequently purified by immobilized metal ion affinity chromatography using an ÄKTAprime plus and Nickel-charged affinity columns (both GE Healthcare Lifesciences, Chicago, USA). Main steps included affinity chromatography with a linear gradient of elution buffer (50 mM Tris-HCl pH=7.5, 150 mM NaCl, 400 mM Imidazole), cleavage of the $His_6$-SUMO tag with SUMO protease (1:50 molar ratio protease to peptide) with simultaneous dialysis (50 mM Tris-HCl pH=7.5, 150 mM NaCl) over night at 4°C, a second affinity chromatography to remove the $His_6$-SUMO tag and finally a size exclusion chromatography with the Superdex 30 16/60 (GE Healthcare) and storage in 1X PBS. The purified protein fragment was used for polyclonal antibody generation and subsequent affinity purification of the antibody (Kaneka Eurogentec S.A., Belgium).

To exclude possible off-targets of the antibody and to validate whether the protein was correctly detected by the antibody (23), we performed a combination of *Tc-rx* in situ hybridisation (DIG-labelled full length probe, 0.4 µl in 30 µl hybridisation buffer) and Tc-Rx antibody staining in *Tribolium* embryos (Fig. S1)(24,25). We found a high degree of overlap between the antibody staining and in situ hybridisation (Fig. S1). No additional staining in the embryo was observed, so that off-targets seem unlikely. To confirm specificity for the endogenous protein, we performed parental RNAi against *Tc-rx* (1.5 µg/µl) following standard procedures (24). We then performed antibody stainings, including a control staining against Engrailed (to exclude differences in staining intensity) in knockdown and wildtype animals (Fig. S1). All steps from fixation to imaging were performed using a standardized protocol. Maximum intensity projections of 34 animals were grouped into three different Tc-Rx staining intensity groups. A blinded categorisation into wildtype and knockdown animals was performed and revealed that all knockdown animals belonged to middle or low strength categories confirming a reduction of Tc-Rx. Hence, the new antibody against Tc-Rx is highly specific for the provided antigen (affinity purification) and the endogenous protein (Fig. S1).

**KNE001 cloning and map**

The vector KNE001 (Vector map S1, pET SUMO-GoldenGate) was based on pET SUMOadapt (modified from the pET SUMO expression vector; (26–28); material transfer agreement with Cornell University, U.S.A., ThermoFisher Scientific, MA, USA). This vector contained an adapter sequence with most importantly a *Bsa*I type IIS recognition site, allowing residue-free cloning of the CDS of interest in-frame with the *ATG::6xHis::SUMO* open reading frame. A second *Bsa*I site was then integrated by first amplifying a fragment additionally containing lac promoter, *CAT* gene and *ccdB* death cassette (29,30) with primers *GG_ccdB_F* and *GG_ccdB_R* (containing a *Xho*I-site) from pTALEN(NI)v2 (gift from Feng Zhang, Addgene Plasmid # 32189, (31)). Second, a *Not*I/*Xho*I digestion resulted in a 1.5 kb *Not*I_*lacP*-

*CAT_ccdB_Xho*I fragment, which was ligated into the *Not*I/*Xho*I linearized pET SUMOadapt. Third, the new pET SUMO-GoldenGate was transformed in ccdB Survival™ 2 T1R Competent Cells (ThermoFisher Scientific, MA, USA).

Hence, by adding GoldenGate linker sequences (S2 Table) that contain *Bsa*I cleavage sites (which do not equal the enzyme's recognition site) to the gene-specific forward and reverse primer, pET SUMO-GoldenGate and the CDS - in our case the N-terminal part of Tc-Rx - can be cut and ligated into a product lacking the original restriction sites. The ccdB cassette in the original KNE001 vector facilitates selection of clones with the gene fragment incorporated.

**Generation of a *Drosophila* bicistronic Rx transgenic line**

In order to generate a comprehensive picture of projections of all Dm-Rx-positive cells and to enable subsequent comparative development of Rx-positive cell groups, we generated a bicistronic line (Fig. S3) using the CRISPR/Cas9 technique (6,32). We also screened available transgenic lines, i.e. two VT-GAL4 lines (https://stockcenter.vdrc.at) that include small fragments of the Dm-Rx regulatory region and hence only covered very small portions of Dm-Rx expression (data not shown).

We built a bicistronic construct as part of the CRISPR repair template, consisting of the C-terminal part of the *Dm-rx* gene, the CDS encoding for *EGFP* and a *P2A* peptide sequence (21,33). The 22 amino acid long P2A peptide (21) is suggested to cause ribosomal skipping (34). This sequence, if placed between two genes or CDS enables the transcription of one long mRNA of *Dm-rx-P2A-EGFP*, but the translation of two separate proteins. The *P2A* and *EGFP* sequences were inserted by using a guide RNA with the target sequence near the *Dm-rx* STOP codon (guide A, Fig. S3). This should result in a common expression of Dm-Rx and EGFP in the same cells, without disturbing the function of either gene through e.g., a fusion product, but with EGFP being in the cytoplasm and Dm-Rx retaining its nuclear localisation.

We included the fluorescent eye marker *3XP3-DsRed* (35). Note that we avoided other eye or body markers, such as *mini-white* because of their size, which might reduce rates of homology-directed repair further. In order to reduce the possible influence of the 3XP3 promotor on Dm-Rx or EGFP expression we inserted the eye marker in the downstream intergenic region, by using a gRNA targeting the intergenic region (guide B, Fig. S3). To facilitate homology-directed repair we included two flanking homology arms (Fig. S3, Vector map S2). As a result, our repair template consisted of seven fragments, which we assembled using a Gibson Assembly® kit (New England Biolabs, MA, USA), following the manufacturer's instructions:

1. Backbone: pJET 1.2/blunt (K1231, ThermoFisher Scientific, MA, USA), EcoRV linearized
2. left homology arm (F1 (Fragment 1): 1 kb of the C-terminus of *Dm-rx* (CG10052) excluding STOP codon
3. P2A peptide (F2): insect codon-optimized sequence (Vector map S2) from plasmid KNE020 (unpublished)
4. EGFP (F3): from plasmid gifted by the Wimmer department, University of Göttingen
5. 3' UTR and intergenic region from genomic DNA (F4): is the region between guide A and B3 (Fig. S3)
6. 3XP3-dsRED-SV40 (F5): eye marker, from plasmid gifted by the Wimmer department, University of Göttingen
7. Right homology arm (F6): 1 kb downstream of guide B3 cut site (three base pairs upstream of its PAM)

The target for guide A would thus be between F1 and F2, and the target for guide B between F5 and F6.

In order to identify single nucleotide polymorphisms in the target strain, integrate the right sequences for F1, F4, F6 and to identify suitable target sites and guide RNAs, we isolated genomic DNA (as described in (6)) from the *Act5C-Cas9, DNAlig4[169]* donor stock (7), and PCR amplified and sequenced the *Dm-rx* C-terminal region, 3' UTR and intergenic region (primers *DmRx_CDS_3'UTR_fw, DmRx_CDS_3'UTR_rev, DmRx_3'UTR_int-region_fw, DmRx_3'UTR_int-region_rev*, Table S2). These regions were used to locate target sites (Fig. S3A[iii]) via the CRISPR Optimal Target Finder (http://targetfinder.flycrispr.neuro.brown.edu/). No off-targets were present for all targets selected. Annealed oligonucleotides were cloned into a U6:3-*Bbs*I vector (based on pCFD3-dU6:3gRNA, Addgene #49410, (36), kindly provided by Hassan M.M. Ahmed (Wimmer department, University of Göttingen, unpublished)) via a GoldenGate reaction, following procedures in (6) but using *Bbs*I

(New England Biolabs, MA, USA). Successful cloning was verified by sequencing the complete chimeric RNA scaffold (including trans-activating crRNA, (36)). guideRNAs were quality controlled by using a T7 Endonuclease I assay (see (6) for procedure). Injection procedures followed descriptions in (37).

Based on the T7 Endonuclease I assay, we selected one guide for the guide A target site, and three with overlapping target sites for the target of guide B (B1-3) (Fig. S3A$^{iii}$).

Next, we designed a 1 kb long F1 (left homology arm) so that it ended before the STOP of *Dm-rx*, as Guide A caused a Cas9 cut only 8 bp downstream of the *Dm-rx* STOP. F4 was designed from the 3'UTR start to the cut site of guide B3 (note that guide B1 and B2 were near B3), with modifications of all PAMs in primers P7 and P8. F6 was 1 kb long, starting at the cut site of guide B3.

All fragments for the Gibson Assembly® were amplified using the primers P1 to P12 (Table S2), containing appropriate overlaps to the neighbouring fragment. F1, 4 and 6 were amplified from the previously isolated genomic DNA of *Act5C-Cas9, DNAlig4* line [169]. We then used three assembly reactions (roman numerals in primers P1 to P12). The first assembled F1 to F3, the second F4 to F6, and the third assembled the products of the first two reactions.

The four plasmids containing each guide and MF01 were precipitated (6,37) to ensure DNA purity and increase viability of embryos after injection. We then made three injection mixes, each containing one of the guides B1 to B3 (250 ng/µl), guide A (250 ng/µl) and MF01 (400 ng/µl), diluted in 1x injection buffer (37). Subsequent injections followed descriptions in (37).

We injected 1203 embryos of which 424 $G_0$ adults survived. We crossed them singly to three $w^{1118}$ virgins of the opposite sex of which 224 $G_1$ crossings gave rise to offspring. We then screened them under a fluorescence stereo microscope (Leica M205 FA, Leica, Wetzlar, Germany) for the presence of the 3XP3-DsRed eye marker and identified 27 positive independent lines. Note, however, that we observed heritable variability in strength and location of DsRed inside the *Drosophila* eye. We thus took four of the 27 positive stocks, with varying degree of eye marker strength and screened wandering third instar larval brains for any detectable differences in the presence of a GFP fluorescence signal resembling known Dm-Rx antibody staining (1). All four stocks did not vary in GFP expression and showed equal similarity to a Dm-Rx antibody staining. To verify this tendency, we performed immunostainings in offspring embryos of these four lines and detected GFP and Dm-Rx signal through a GFP antibody staining. Embryos of all four stocks showed near 100 % overlap to Dm-Rx and a cytoplasmic signal. Finally, to verify that insertion was performed as planned (Fig. S3A$^{ii}$), we isolated genomic DNA from one whole adult male of each of the four stocks using the Zymo Research *Quick*-DNA Miniprep Plus kit (Zimo Research, Irvine, CA, USA) following the manufacturer's Solid Tissues Protocol. We then amplified DNA fragments containing the whole region by nested PCR (primers *DmRx_trans-ver_fw, DmRx_trans-ver_rev, DmRx_trans-ver_nested_fw, DmRx_trans-ver_nested_rev*, Table S2). We sequenced the regions surrounding the cut sites with primers *DmRx_trans_seq_Ct_fw* and *DmRx_trans_seq_iRe_rev* (Table S2). All four stocks showed correct sequencing at guide A cut sites, but only the line used in this study showed completely correct sequences, thus allowing us to perform suitable experiments and closer characterisation (Fig. S3C-F).

To verify that EGFP is indeed localised in the cytoplasm, we performed immunostainings for Dm-Rx and GFP in embryos. With higher magnification we were able to see a substantial amount of GFP in the cytoplasm surrounding the nuclei marked by Dm-Rx (Fig. S3C) and DAPI (not shown). We also wanted to know whether expression from the transgenic Dm-Rx locus was qualitatively different from the endogenous expression, so to ensure that we investigated Dm-Rx expression similar to a wildtype situation. For this we performed immunostainings against Dm-Rx in the adult *Drosophila* brain with identical settings and imaged them identically (Fig. S3D). We were not able to detect any absence of domains (Fig. S3D). Differences between the wildtype strain $w^{1118}$ and our transgenic line were – if present – as large as differences between individual brains of the same genetic background.

We then tested whether the expression of Dm-Rx and EGFP in the same cells is maintained in the adult brain (Fig. S3E-F). Indeed, by qualitative assessment we were able to see an approximately 100 % coexpression, with prominent projections marked as well (Fig. 3).

We thus concluded that the Rx-GFP bicistronic line was suited for our use.

**Characterisation and validation of the *Tribolium* Rx-GFP enhancer trap**

We identified a suitable *Tribolium* transgenic line in the GEKU base website (# E01101, http://www.geku-base.uni-goettingen.de/; (4)). Insertion had been mapped to the upstream region of *Tc-rx* ((4); Fig. S2A). To identify to which degree Tc-Rx expressing cells also express GFP we performed co-stainings in adult brains. We found that n-ventral Tc-Rx-positive cells were not marked by the line at all while n-dorsal domains were only partially marked (Fig. S4). However, by manually checking each GFP expressing cell, we found that all cells expressing GFP in the region surrounding the protocerebral bridge, also expressed Tc-Rx (Fig. S2B-D). Hence, there were no cells that were marked false-positively. Interestingly, there were more GFP expressing cells showing overlap to Tc-Rx expression in all other stages of development (Fig. S2E). To ensure that Tc-Rx is expressed similar to the wildtype situation, we performed identical immunostainings against Tc-Rx and imaging with identical settings in the transgenic line and wildtype $v^w$ background (Fig. S2C)(38). We found that differences between conditions were no larger than the differences observed between individuals of the same condition. We thus concluded that the Rx-GFP enhancer trap was suitable for further experiments.

**Generation of homozygous stocks of Rx-GFP transgenic lines**

A homozygous stock of the *Tribolium* Rx-GFP enhancer trap was generated by genotyping adult wing tissue, as described in (6,39), with primers *GEKU-Rx-GFP_wt_fw*, *GEKU-Rx-GFP_wt_rev*, *GEKU-Rx-GFP_trans_rev* (Table S2).

A homozygous stock of the *Drosophila* Rx-GFP bicistronic line was generated by crossing the male offspring ($G_2$) of the $G_1$ cross to female virgins of a $w^-$; $wg^{Gla-1}/CyO$ $2^{nd}$ chromosome balancer (a kind gift by the Wimmer department, University of Göttingen). CyO/$w^-$ positive animals (G3) were selected and crossed to each other, to create homozygous positive animals (G4) for the transformation marker (*3XP3-dsRed-SV40*).

Both transgenic lines were viable in the homozygous background.

#### R45F08-GAL4 crosses

To reveal the overlap of secondary cells of the DM1-3 and 6 lineages marked by the R45F08-GAL4 line (12,13) with Dm-Rx expressing cells we performed two crosses and subsequent immunostainings (Fig. S5).

First, we crossed the R45F08-GAL4 line with a UAS-mcD8::GFP line and investigated offspring third instar larvae to visualize the characterized cells and subsequently stained with anti-GFP and anti-Rx antibodies (Fig. S5A-B).

Second, to visualize an overlap of Dm-Rx expressing cells and R45F08 labelled cells, we first crossed the *Drosophila* Rx-GFP bicistronic line each separately with R45F08-GAL4 and UAS-mcD8::RFP. The respective offspring was then crossed to each other. We then dissected 15 brains of third instar larvae, stained them with anti-RFP and anti-GFP, screened for the presence of RFP and GFP label and imaged double-positive brains (Fig. S5C).

#### Staging of *Tribolium* and *Drosophila* animals

Table S6 displays all stages and their description included in our study. Particularly pupal staging and the late larval stages were determined using time (which allowed us to calculate relative times of pupation) and morphology as criteria to confirm the timed staging.

*Drosophila* embryonic stages were determined using the staging of (14) and pupal stages using staging in (15) (Table S6 displays the most important pupal selection criteria). Eye colouring and morphology were not included in staging due to the $w^{1118}$ background. Information on the length of embryonic development used in Fig. 10 was derived from (14). Length of larval development and pupation was measured for our Rx-GFP bicistronic line specifically.

*Tribolium* embryonic stages were determined using the staging of (3) and for late embryonic stages using staging of Scholten and Klingler (unpublished). *Tribolium* pupal and late larval staging was aided by (40) and Dippel (unpublished). Information on length of embryonic development used in Fig. 10 was derived from (3) and Scholten and Klingler (unpublished). Total developmental time was taken from (41). Larval and pupal developmental length was measured for our Rx-GFP enhancer trap specifically. To that end, we first determined the duration of pupation in the used Rx-specific transgenic lines. Pupation in the *Tribolium* Rx-GFP line took approximately 140 h, while pupation in the *Drosophila* Rx-GFP line took approximately 100 h. A developmental progress of 5 % equals 7 h in *Tribolium*, and 5 h in *Drosophila*.

**Specimen fixation and immunohistochemistry**

Methanol fixation of *Drosophila* embryos was performed following standard protocols (42). Fixation of *Tribolium* embryos was based on (43) with following modifications: Fixation was performed with 2 ml of fixation buffer PEMS (0.1 M PIPES, 2 mM MgCl, 5 mM EGTA, pH = 6.9); we added 180 µl of 37 % formaldehyde (F 1635, Merck, Darmstadt, Germany) and fixed embryos between 25 and 32 min; devitellinisation was first conducted with a 0.9 µm canula, for older stages (> 40 h) we followed with a 0.8 µm canula.

Immunohistochemistry of embryos was based on procedures in (25), with the addition of preceding washes in a descending methanol series (75, 50 and 25 % Methanol with PBS-T 0.1 %), followed by two rinse steps and three 10 min washes.

For all stainings normal goat serum was used as blocking solution (NGS, G9023, Merck, Darmstadt Germany, see Table S3). Fixative for all other stages except for embryos was 4 % PFA (wt/vol, paraformaldehyde, (e.g. P6148, Merck, Darmstadt, Germany) in PBS, 130 mM NaCl, 7 mM $Na_2HPO_4$, 3 mM $KH_2PO_4$, (44)). Washing buffer for all stages except embryos was phosphate buffer (PB, see (20) for recipe). Brains were dissected using Dumont No. 5 forceps in ice-cold PB. All steps were performed in 180 µl volume in 9-well PYREX™ Spot Plates (ThermoFisher Scientific, MA, USA) on an orbital shaker. Protocols were adapted from (20,44).

**Image acquisition, processing and 3D reconstruction**

If not otherwise specified, imaging was performed at a Leica SP8 confocal microscope (Wetzlar, Germany). Objectives used were either a Leica apochromat 20x (NA = 0.75) or a 63x HC PL APO CS2 (NA = 1.30) glycerol-immersion objective. DAPI was excited by a Diode laser (405 nm), Alexafluor 488 (ThermoFisher Scientific, MA, USA) by an Argon laser (488 nm), Alexafluor 555 by a DPSS laser (561 nm) and

Alexafluor 647 by a HeNe laser (633 nm). Detection was performed with Hybrid detectors and photomultipliers, at an 8-bit depth. Averaging was depending on which staining was performed, set on line or frame averaging of 4. Step size was set to system optimized values defined by the LASX software. Image size was set between 1,024 x 1,024 and 2,048 x 2,048 pixels. Images were processed, adjusted for brightness and contrast, cropped, merged and rotated using the Fiji software (45). Maximum intensity projections and smooth manifold extractions (SMEs; (8)) to retain 3D spatial relationships were calculated using Fiji as well (45).

3D reconstructions were performed in Amira 5.4.1 (Visage Imaging, Fürth, Germany). We created Labelfield data with the same pixel and voxel size resolution as the original data set. We then used the Segmentation Editor to identify and create material for each tract and central complex neuropils by employing the Wand tool. Subsequent marking was modified for visual ease using the grow, fill holes and smooth functions of the Segmentation Editor. Subsequently we created 3D surfaces with the Surface Generator.

In some cases, projections were too thin to be recapitulated in the 3D surface. For this, where we logically inferred a connection of axons that was only faintly marked by the original file, we used the Brush tool.

We only reconstructed the axon connections to certain cell bodies where we were sure that they are directly connected. This excluded a few cell bodies from the analysis, particularly in the *Drosophila* adult brain.
